# Repeatability of an extended phenotype: potential causes and consequences of nest variation in the Northern House Wren (Troglodytes aedon aedon)

**DOI:** 10.1101/2024.03.08.584149

**Authors:** Chandler E. G. Carr, Zoë M. Swanson, Dustin G. Reichard

**Author notes:** Corresponding author: Chandler E. G. Carr.

## Abstract

**LAY SUMMARY:**

- Repeatability of construction behaviors in the wild is understudied
- House Wrens are a cavity-nesting songbird
- House Wren nests are highly variable in their morphology
- Female House Wrens are repeatable in cup composition, but not nest dimensions
- The dimensions of the nest are built to match the dimensions of the cavity
- House Wren nests are unrelated to nestling survival or female condition
- Female preferences for nest building may explain nest variation

Construction behavior is an aspect of the extended phenotype that allows organisms to build structures that alter their environments in potentially beneficial ways. Although individuals vary in the expression of this extended phenotype (e.g., structure morphology), the repeatability of construction behaviors remain understudied, especially among free-living populations. Many oviparous taxa construct nests, making them of particular interest because variation in nest architecture may directly affect fitness. Using a free-living, cavity-nesting songbird, the Northern House Wren (*Troglodytes aedon aedon*), as our model, we estimated the contribution of the primary builder (the female) to nest variability by measuring the repeatability of nest morphology between successive clutches. We further examined whether nest morphology was related to the dimensions of the nesting cavity, breeding date, or nest success. We found the composition of the cup lining to be a highly repeatable behavior for the nesting female, although the size and composition of the structural platform appeared more related to the dimensions of the cavity. Nest morphology remained variable throughout the breeding season, showing no significant correlations with breeding date, and it was unrelated to clutch size or offspring survival. Our study suggests that variation in construction behavior is likely the product of multiple factors including the preferences of the builder and physical constraints. The absence of any clear links between construction behavior and fitness indicates that nest morphology is not under strong selection. As a result, diverse female building preferences may explain the extreme among-individual variation in nest structure in this species.

## INTRODUCTION

Alfred Russel Wallace (1865) once stated: “…so great is the tendency to vary that it is difficult to find two individuals exactly alike.” Phenotypic variation is both ubiquitous and a fundamental requirement for evolution (Darwin & Wallace 1858). As such, a considerable amount of research has been devoted to understanding the causes and consequences of among-individual variation. The interaction between an organism’s phenotype and the environment determines its success, and selection can push populations toward one or more optimal phenotypes (van Rijssel et al. 2018, LeGrice et al. 2019). However, the relationship between the phenotype and the environment can be bi-directional, allowing individuals to reshape their environments in ways that affect their fitness (i.e., “niche construction”; Laland et al. 1999). For example, an individual managing a thermoregulatory challenge might rely on costly physiological mechanisms like shivering or increasing adipose tissue, but an alternative behavioral mechanism, such as constructing a burrow, could insulate the animal through the construction of a *new* environment, thereby buffering against this selective pressure (Clark et al. 2020).

Construction behavior is a unique behavioral phenotype where the product of the construction is an extension of the builder (i.e., “extended phenotype”; Dawkins 1982) that modifies the environment to be more beneficial for the builder. Construction behaviors may enhance the builder’s fitness by providing protection from predators (Beebe 1953, Hermes, 2013, Sugiura 2016) and environmental exposure (Zeibis et al. 1996, Korb & Linsenmair 2000, Korb 2003, Rosell & Campbell-Palmer 2022), as well as increasing access to resources such as food (Hölldobler & Wilson 1990, Miyashita & Shinkai 1995, Blamires et al. 2010) and mates (Borgia 1985, McKaye et al. 1990). As a result, construction behavior has become taxonomically widespread across both vertebrate and invertebrate species (Hansell 2005). Eusocial insects build colonial shelters (e.g. termite mounds), some mammals dig burrows, most spiders spin webbed traps, and certain species of fishes and birds construct bowers.

However, despite the clear potential for construction behavior to benefit the fitness of an individual, trade-offs often occur between investment in the construction process and other behaviors that affect fitness which can result in negative effects on both the builder (Madden 2002, Bailey et al. 2015) and the structure (Ellendula et al. 2021, Parthasarathy et al. 2022). For example, investing more time in the construction process potentially reduces available energy for foraging or mate-searching.

Many organisms build multiple versions of the same structure throughout their lifetimes, which provides an opportunity to evaluate the consistency of this behavior. It is well established that the products of construction behaviors exhibit among-individual variation, but the extent and cause of variation *within* an individual’s construction behavior is less clear (DiRienzo & Aonuma 2018). For example, an individual’s construction behavior might be consistent, plastic, or some combination of the two as its experience and environment changes. Repeatability estimates, calculated as intra-class correlation coefficients, can be used to assess how much of the variation observed among individuals can be explained by variation within individuals (Hayes & Jenkins 1997). When among-individual variation is high and within-individual variation is low, then the repeatability estimate will be high and that trait will be considered repeatable. Repeatability estimation can be used to understand the extent to which phenotypic traits are the product of static characteristics, such as heritability (“upper level of heritability”; Bell et al. 2009, Møller 2006) and individual preferences (Mennerat et al. 2009, Järvinen & Brommer 2020, Silva et al. 2024), or plastic characteristics, such as experience (DiRienzo et al. 2019, Whittaker et al. 2023). For this reason, the repeatability of a large variety of behaviors have been examined across taxa (Bell et al. 2009), but there are few studies examining the repeatability of construction behaviors. Most of these studies have focused on spider web structure (Fisher et al. 2019, DiRienzo et al. 2020, DiRienzo & Montiglio 2016) or nest morphology (Rushbrook et al. 2008, Muth & Healy 2011, Whittaker et al. 2023), but few have examined the behavior exclusively in the wild (Møller 2006, Järvinen et al. 2017, Silva et al. 2024).

Nest building is a widespread construction behavior among oviparous taxa that serves the primary function of providing a safe, hospitable space for eggs or altricial young to develop (Mainwaring et al. 2014a). With few exceptions (e.g., brood parasites), bird species produce an incredible array of nests that can be structurally complex or simple, but must contain at least one of four zones: an outer nest layer, an attachment to the environment, a structural layer, and a lining (Hansell 2000). For example, scrape nests, built on the ground, may be considered just structural while nests built in trees often require an attachment, structure, and a cup lining.

Zones are often discernible from one another by the materials used, with zone composition being largely species specific (Biddle et al. 2018). Nests can be built by the male, female, or both with some division of labor by zone (Hansell 2000). The morphology of the nest, structure, size, and composition, can be important for providing the same benefits of niche construction observed for other structures by protecting the young from predators (Møller 1990, Hays et al. 2022) and environmental exposure (Franklin 1995, Mainwaring et al. 2014b, Cerezo & Deeming 2016, Perez et al. 2020, Medina et al. 2022), as well as potentially playing a role in mate attraction for some species (Ueda 1985, Mainwaring et al. 2008, Jelínek et al. 2016).

Here, we investigated potential causes and consequences of the variation and repeatability of Northern House Wren (*Troglodytes aedon aedon*) nest morphology and composition. House Wrens are secondary cavity nesters that raise two to three clutches a season, often reusing the same nest throughout (Johnson 2020). They will, however, build a new nest when the previous nest is stolen, destroyed, or experimentally removed (Alworth 1996), making them an ideal model for the examination of repeatability in construction behavior. House Wren nests contain 2 to 3 zones: sometimes an outer layer consisting of spider egg sac material, a structural zone made of various types of sticks, henceforth referred to as the “stick platform,” and a cup defined as a simple depression, at the top of the stick platform, lined with grasses, feathers, and other material (McCabe 1965). Although the layout and composition of the zones are considered standard, House Wren nests are highly variable in their morphology. For example, in some nests the stick platform may be built so large that it covers the entrance hole of the box (McCabe 1965; Figure S1A) while other nests have a stick platform only a few centimeters in height (Figure S1B). While establishing a territory, unmated males place a variable number of sticks (range: <10 to >400; Johnson 2020) into potential nesting cavities. After pairing, the female completes the stick platform (Alworth & Scheiber 2000, Johnson 2020) and creates a lined cup either before or during egg laying. Considering the extensive effort of the female in both completing the stick platform and constructing the nest cup, we defined the female as the primary nest builder in our study.

Although the architecture of the nests of other bird species have shown to be beneficial to the fitness of the builder (Mainwaring et al. 2014), House Wren nest architecture appears to be unrelated to the mate selection process, the survivability of offspring (Alworth 1996, Eckerle & Thompson 2005), or the limiting of brood parasitism (Pribil & Picman 1997). Thus, the available evidence suggests that nest architecture in this species has limited effects on fitness, and so it remains unclear why House Wren nests exhibit such extreme variation among individuals. The absence of selective pressure on nest architecture could be one explanation for the amount of variability observed. Alternatively, some research suggests that constraints imposed by the cavity dimensions and the accessibility of building materials may also drive variation in nest architecture (Stanback et al., 2013). However, no study to date has explored the role of the preferences of females during nest construction, such as the types and quantity of material used, as an explanation for the variation. By sampling successive nests built by the same female one can investigate whether nest architecture is repeatable at the individual level. If nest architecture *is* repeatable, then differences between individual females in their building preferences can provide a previously underappreciated explanation for the high intraspecific variation in House Wren nests.

We sampled House Wren nests across an entire breeding season to assess the contribution of the female builder, the box cavity, and breeding date to nest architecture. We also examined whether variation in nest characteristics correlated with one measure of female body condition (residual between the individual’s mass and tarsus length), clutch size, or offspring survival, which provided proxies of fitness. Unlike most construction behavior research that often relies on coarse measures of structure like nest weight (Mainwaring et al. 2008, Walsh et al. 2010, Adreani et al. 2022), we used a finer approach of directly quantifying the individual building materials (e.g., exact count of sticks) used in nest construction.

We predicted that nests would have high variability among individuals but that there would be low within-individual variability. Specifically, since the female builds the majority of the structure after pairing, the female identity would be the best predictor of nest dimensions and composition as measured by a repeatability estimate. If supported, this outcome would highlight the importance of female preferences for specific nest characteristics and building materials in determining nest architecture. Furthermore, based on previous research in other House Wren populations (Alworth 1996, Eckerle & Thompson 2005, Stanback et al., 2013), we predicted that nest architecture would not relate to offspring survival, but may relate to the dimensions of the cavity in which it was built or the breeding date when it was measured.

Successive nests that are built in the same box should exhibit repeatability for those characteristics that are most affected by the box’s dimensions. Furthermore, seasonal changes in other variables such as weather and vegetative cover might favor different nest architecture as breeding progresses. For example, thinner nest cups could be favored to provide less insulation as average temperatures increase (Mertens 1977). Alternatively, because clutch size declines in this species across the breeding season (Johnson 2020), females might build smaller nests over time due to fewer offspring or a more general decline in reproductive investment later in the season.

## METHODS

### Study System

We monitored Northern House Wren (*Troglodytes aedon aedon*) breeding behavior in 160 nest boxes across six sites in Delaware and Union counties in Ohio, U.S.A., between May 9 and August 25 of 2022. Of those 160 boxes, 79 produced at least one nest with a complete clutch, which was the threshold for inclusion in our study. With the exception of three nest boxes located in suburban yards, the majority of nest boxes were located in edge habit separating grass from forest. We measured nest box dimensions with digital calipers (nearest 0.001 mm). Nest boxes were rectangular wooden cuboids with internal dimensions of approximately 110 x 115 x 195 mm (Figure 1, see Table 1 for variation in box dimensions). Each box possessed a circular entrance hole with an average diameter of 34 mm and depth of 46 mm, drilled an average of 119 mm from the bottom of the box. Most (*n* = 73/79; 92.4%) boxes also had a rectangular gap separating the top of the front or side panel from the roof that allowed the panel to swing open. This gap was the same width as the box with a height ranging from 4 mm to 30 mm. Some wrens used this gap as an entrance, particularly when their nest obstructed the primary entrance hole. The variation in box dimensions were due to imperfect construction and the preferences of landowners for slightly different box designs. All boxes were suspended about 1.75 meters off the ground by a greased metal conduit pole to limit predation.

**Figure 1:**
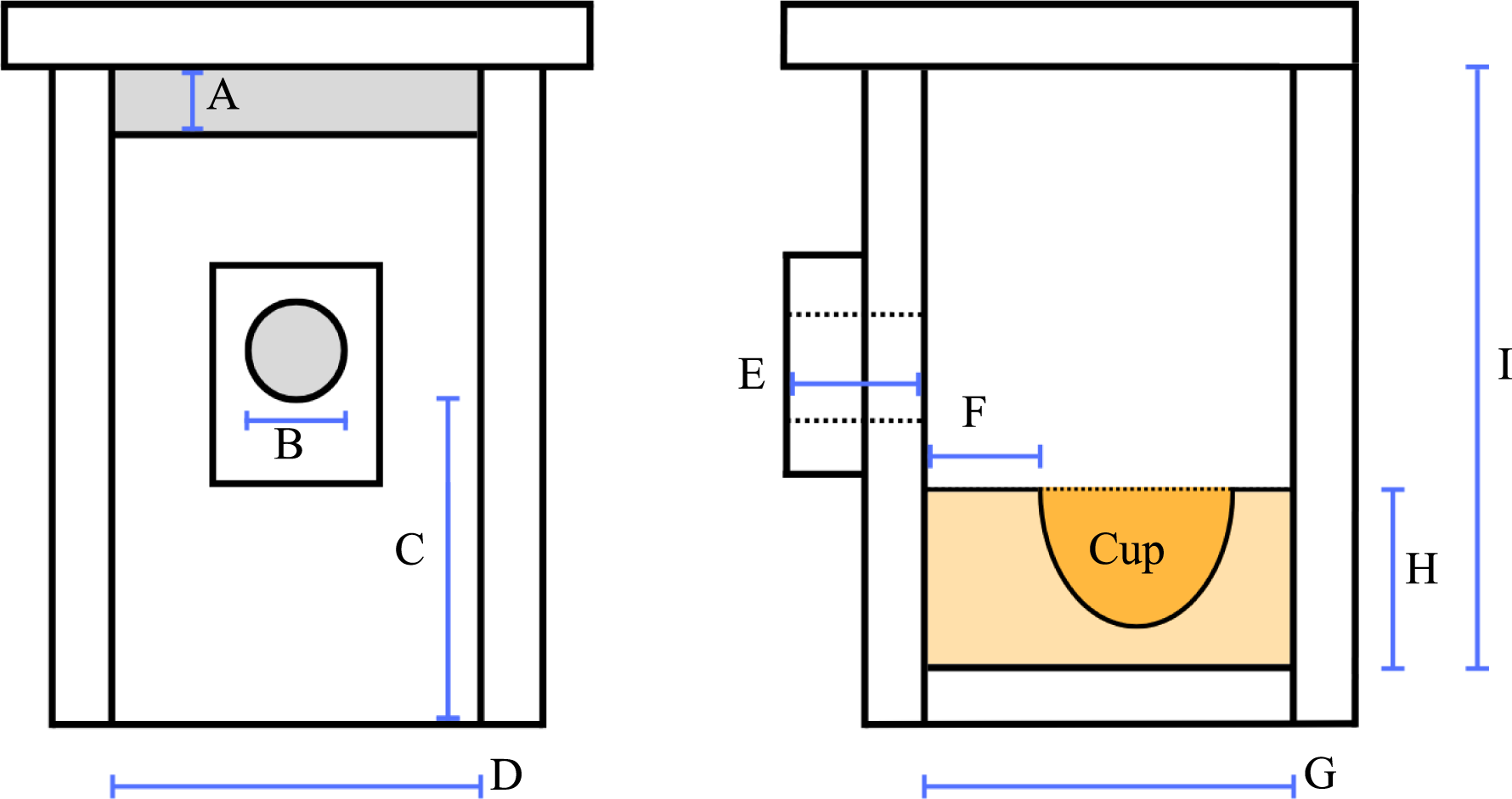
Model of the nest box and nest (in orange). Both the front (left) and side (right) of the box are shown. A: height of the rectangular gap, B: diameter of the entrance hole, C: distance from the bottom of the box to the bottom of the entrance hole, D: internal length of the box and length of the nest, E: depth of the entrance hole, F: width of the berm, G: internal width of the box and width of the nest, H: height of the nest, I: internal height of the box.

**Table 1:** Average, standard deviation, and range of the dimensions of the nest boxes used in this study. See Figure 1 for a model of an average nest box.

|  | Average (mm) | Standard Deviation (mm) | Range (mm) |
| --- | --- | --- | --- |
| Internal Box Height | 195.19 | 14.92 | 172.20 - 219.90 |
| Internal Box Volume | 2462.20 | 434.52 | 1812.99 - 4133.73 |
| Entrance Diameter | 33.60 | 5.22 | 27.90 - 44.65 |
| Entrance Depth | 44.11 | 6.78 | 25.00 - 56.40 |
| Height of Entrance | 118.94 | 21.19 | 83.10 - 151.60 |
| Gap Height (when present) | 14.82 | 6.11 | 4.30 - 29.55 |

We monitored 108 total nests (some boxes were used multiple times), but were unable to visit boxes daily, so exact aging of nests was not always possible. However, House Wrens have predictable rates of egg laying and incubation time, so we were able to estimate first egg dates by counting eggs and incubation start dates by backdating from hatch date (Johnson 2020). In cases where we were not present for hatch day, nestling ages and estimated hatch date were evaluated by measuring nestling weights, wing chord length, and feather tract development as described by Brown et al. (2011).

As soon as we discovered a complete clutch, the female was caught in a mist net at the nest box and banded with a USFWS aluminum leg band and a unique combination of plastic colored bands for future identification. It was essential to capture and band females to sample repeated nesting attempts, but we chose not to capture males to minimize further disrupting both our experiment and a separate, unrelated study. The female’s weight (nearest 0.1 g; Pesola spring scale), tarsus length (nearest 1 mm; ruler), and wing length (nearest 1 mm; ruler) were also measured. While the female was processed, we measured the height of the nest and the width of the “berm” (Stanback et al. 2013), which was defined as the front wall of the cup directly against the entrance hole (see Figure 1), using a dial caliper (nearest 0.1 mm). On day eight of nestling life, we collected measurements and banded each nestling as described above for adult females. We continued to monitor nests until the clutch fledged or failed, at which point the nest was collected in a large and resealable plastic bag for dissection in the lab and to prevent the nest from being reused or built upon for a successive clutch.

### Nest Analysis

After collection, we placed nests in a −20 C freezer for 24 - 48 hours to kill any arthropods. For dissection, nests were placed on a large sheet of white paper and carefully pulled apart to quantify nest materials and characteristics. Most nests were dissected by the lead author, but two assistants were also trained for dissection. We made exact counts of individual sticks, grasses, plumulaceous, and pennaceous feathers. Sticks and grasses needed to be larger than an arbitrary length of 4 cm in order to be counted. Smaller pieces of vegetation were not quantified along with other debris not described below. Down, contour, and semiplume feathers were categorized as plumulaceous, while wing and tail feathers were categorized as pennaceous. We used an ordinal scale of 0 to 3 to quantify the amount of fecal material and spider egg sacs present within each nest. Finally, we noted, as a binary yes or no, the presence of snake skin and anthropogenic objects (e.g., plastic).

### Statistical analysis

Although we sampled 108 nests in total, a small number had to be excluded from our final analysis. One nest was removed during dissection because it contained signs of reuse (i.e., two individual grass layers separated by a thick stick layer). Two nests were not collected from the field due to reuse or extensive nestling decomposition after predation, and so were unable to be dissected. Due to data loss, the dissection measurements of two additional nests are unaccounted for. Our final sample size was 103 nests.

R (version 4.2.3) was used for all analyses. We first assessed whether the observed variation in nest measurements was normally distributed using separate Shapiro-Wilk tests. To test whether nest measurements were repeatable within females, we calculated repeatability estimates (packages: rptR, version 0.9.22) with the nest measurement as the dependent variable, and the identity of the nesting female as the grouping factor. We used the functions rptGaussian or rptBinomial depending on the nest measurement being explored. Parametric bootstrapping and permutation, at 10,000 iterations each, were performed to produce confidence intervals and adjusted alpha values for the repeatability estimates, respectively.

Significant repeatability estimates were indicated by 1) confidence intervals not overlapping zero (DiRienzo & Aonuma, 2018) and 2) an adjusted alpha value of 0.05. Where nest measurement was binomial, the link-scale approximation (Stoffel et al. 2017) is reported. We used the same approach to test whether the box identity predicted nest measurements. In these models, the nest measurement was the dependent variable, and the identity of the box was the grouping factor. Although there was some overlap in sampling between models with the same female nesting more than once in the same box, the sample size (5/24; 21% of females reused the same box and 6/27; 22% of boxes were used by the same female) was too small to provide a reliable estimate of the effect of these repeated samples on nest measurements. Thus, we chose to avoid a more complex model that included both box and female identity. Finally, we defined repeatability estimates less than 0.20 as low, between 0.20 and 0.40 as moderate, and greater than 0.4 as high repeatability (Bohn et al. 2017).

Linear regressions (Base R Package) were used to test for a relationship between nest measurements and box proportions, date of measurement, and age of the nest, which was defined as the number of days between the first egg and the nest collection date. We used generalized linear models (Base R Package) to analyze the effect of nest measurements on the outcome of the clutch. These models assumed a binomial distribution with nest outcome (fledge/fail) as the dependent variable. Linear models were used to analyze the effect of nest measurement on clutch size and fledge count, assuming a gaussian distribution. Both the generalized linear models and linear models had the nest measurement as the fixed effect, and the date of nest measurement as a covariate to control for seasonal variation in reproductive output.

## RESULTS

### Variability in Nest Architecture and Correlations Between Nest Components

House Wren nests were highly variable (Figure 2). The ratio of the nest height to internal box height ranged from 0.254 to 0.902 (Figure 2A; 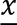 ±SD: 0.59 ± 0.144). Similarly, the volume of the nest ranges from 655.71 mm³ to 2776.96 mm³ (Figure 2B; 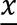 ±SD: 1437.44 ± 409.29). Sticks were the primary contributor to the height and volume of the nest, and we found that stick count was also variable, ranging from 170 to 1491 sticks in a single nest (Figure 2C; 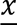 ±SD: 676.14 ± 25.01). The width of the nest berm ranged from 7.00 mm to 77.45 mm (Figure 2D; 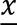 ±SD: 39.62 ± 1.27).

**Figure 2:**
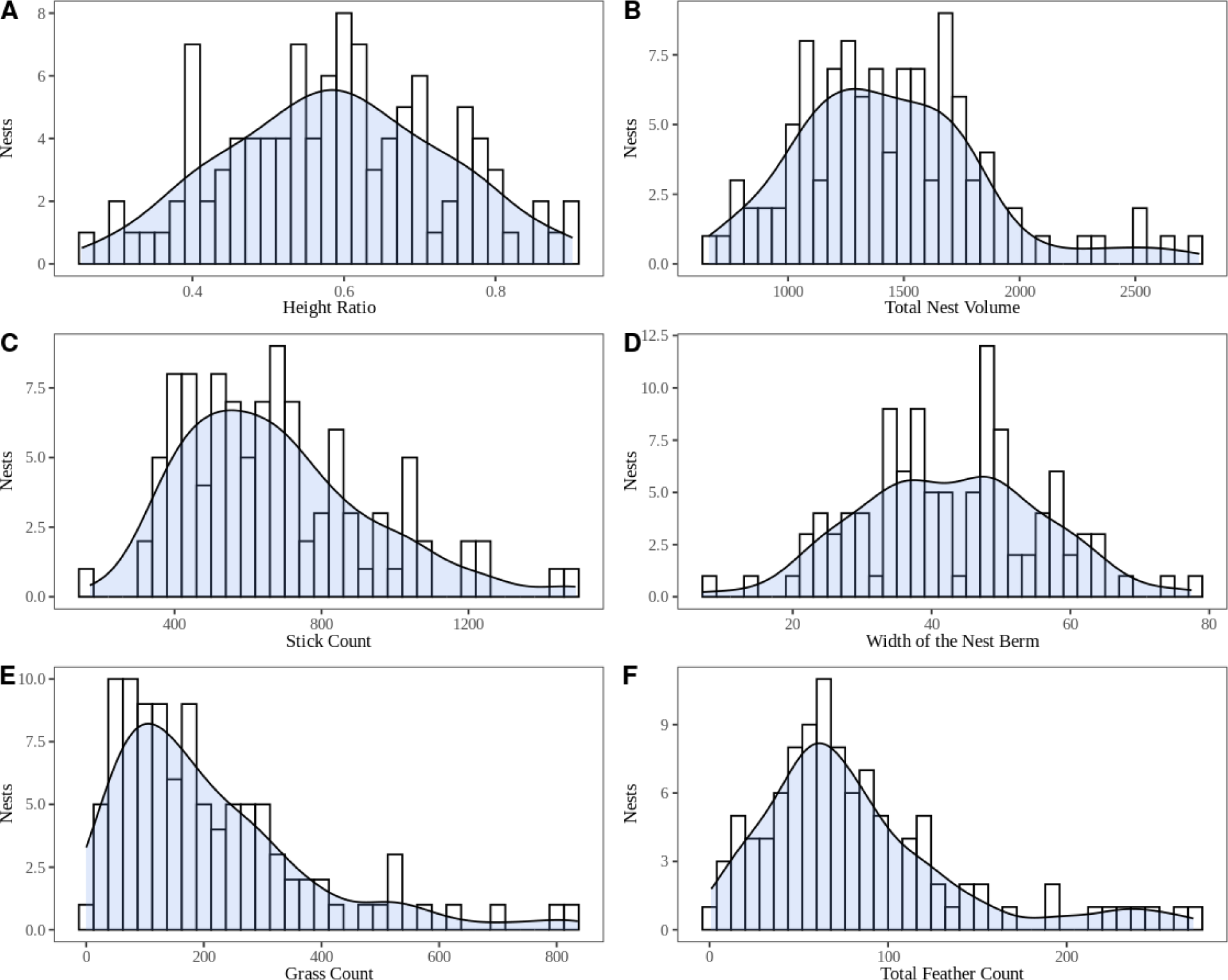
Histograms showing the number of nests that contained the specified variable. “Height Ratio” (A; P = 0.74) and “Width of the Nest Berm” (D; P = 0.94) exhibit a normal distribution, but other factors show a right-skewed distribution (P < 0.001). The blue plots layered over the histograms represent the outcome of an estimated density distribution.

We found the number of grass strands within the cup of a single nest ranged from zero to 836 (Figure 2E; 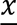 ±SD: 210.19 ± 16.49). Feather count within the cup ranged from a singular pennaceous feather to 271 total feathers (Figure 2F; 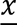 ±SD: 83.79 ± 5.70). Plumulaceous feathers were the most common feather found in nests with an average ratio of 5 plumulaceous feathers for every 1 pennaceous. Although the cup is primarily lined with grasses and feathers, other materials such as snake skin (*n* = 19/103; 18.4%), anthropogenic objects (e.g., plastic; *n* = 54/103; 52.4%), and hair, noted anecdotally but not quantified, from a variety of mammalian species, including human, were observed. We also frequently found varying amounts of spider egg sac material interspersed throughout the stick platform (*n* = 88/103; 85.4%).

The ratio of the nest height to internal box height (P = 0.74; Figure 2A) and the width of the nest berm (Figure 2D; P = 0.94) were normally distributed across the population. All other continuous nest characteristics showed a right-skewed distribution (P < 0.001).

The width of the berm positively correlated with the height (r^2^ = 0.03, *P* = 0.04), volume (r^2^ = 0.09, *P* < 0.001, Figure S2A), and stick count (r^2^ = 0.04, *P* = 0.02, Figure S2B) of the nest. The stick count positively correlated with the height (r^2^ = 0.37, *P* < 0.001, Figure S2C) and volume (r^2^ = 0.41, *P* < 0.001, Figure S2D) of the nest. Total feather count (r^2^ = 0.03, *P* = 0.04, Figure S2E) and pennaceous feather count (r^2^ = 0.06, *P* = 0.005, Figure S2F), but not plumulaceous feather count (*P* = 0.15) positively correlated with the grass count. No other correlations were statistically significant (*P* > 0.19, Table S2).

### Variation Explained by Female Builder Identity and Body Condition

Over the course of the nesting season, 24 females renested in our boxes after their initial clutch. Of those females, 19 renested once and 5 renested twice on our study site, and the majority of those females renested in a different box from any previous clutch (22/29 total renests; 75.9%; Table S1). The identity of the nesting female did not explain the variation in the general nest dimensions, such as the ratio of the nest height to internal box height (*R* = 0.24 [95% CI = 0 - 0.56], *P* = 0.08, Figure 3C), nest volume (*R =* 0.22 [95% CI = 0 - 0.53], *P* = 0.11), or the the width of the berm (*R =* 0 [95% CI = 0 - 0.34], *P* > 0.99, Figure 3E). The female identity also did not explain the number of sticks within the stick platform (*R* = 0.15 [95% CI = 0 - 0.49], *P* = 0.20, Figure 3A). However, when considering the cup lining, the grass material count was highly repeatable for the female (*R* = 0.56 [95% CI = 0.23 - 0.77], *P* = 0.003, Figure 3B). While the total count of feathers in the cup was not repeatable (*R* = 0.04 [95% CI = 0 - 0.39], *P* = 0.37), the pennaceous feather count appeared moderately repeatable (*R* = 0.33 [95% CI = 0 - 0.62], *P* = 0.05, Figure 3F), although we do note that the confidence interval overlaps with zero. The ratio of pennaceous feathers to plumulaceous feathers within the cup of the nest was highly repeatable (*R* = 0.90 [95% CI = 0.79 - 0.95], *P* = 0.002, Figure 3D). No other individual measurement was repeatable (Table 2). No significant correlations between the body condition of the female builder and nest morphology were observed (*P* > 0.11, Table S3).

**Figure 3:**
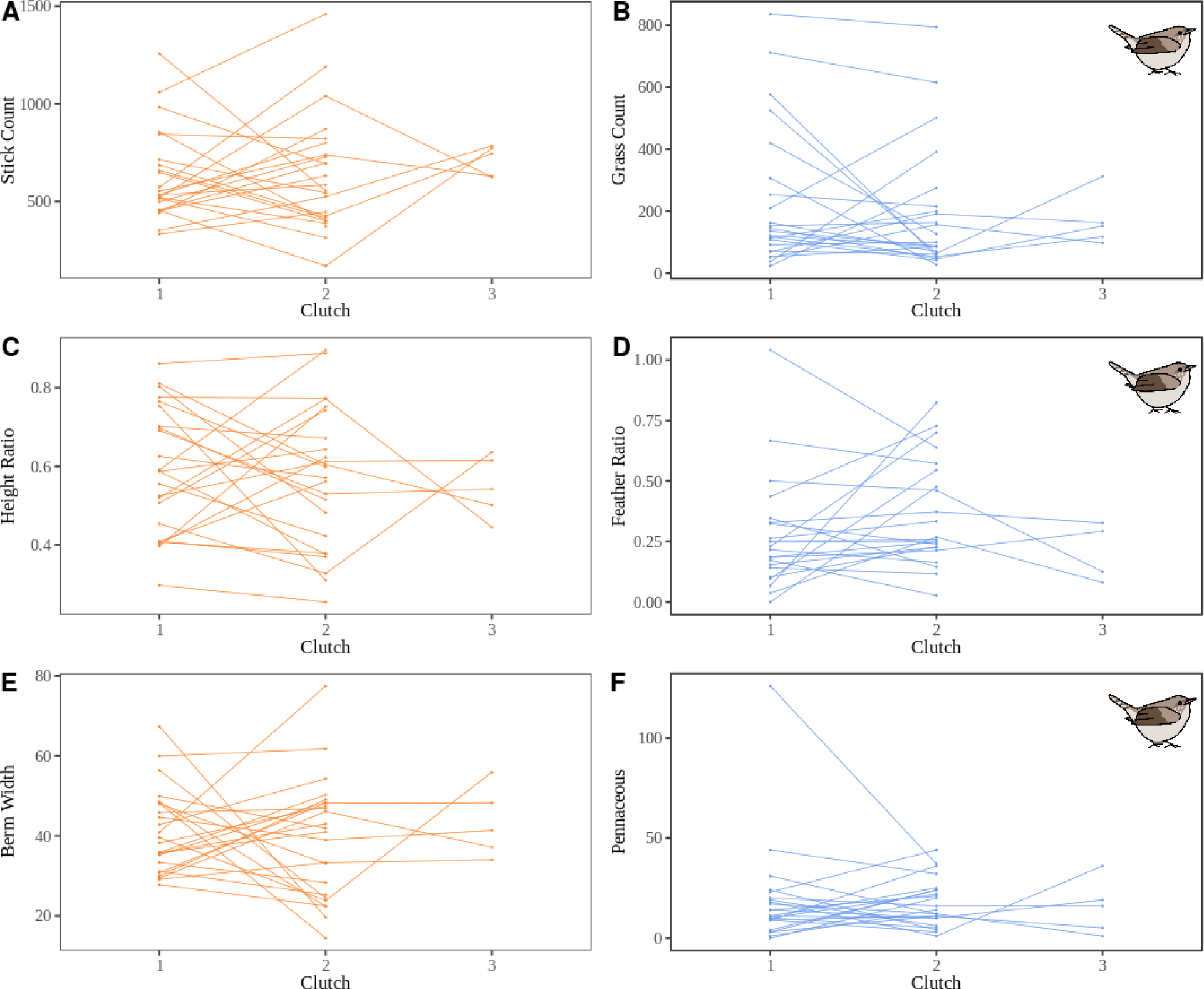
The repeatability of nest measurements across successive clutches for individual females. Characteristics of the stick platform are shown in orange (left), and characteristics of the cup lining are shown in blue (right). Significant repeatability by female identity is indicated by the presence of a House Wren drawing. Female House Wrens construct similar nest cups across multiple nesting attempts with the grass count (B; R = 0.56, P = 0.004), feather ratio (D; R = 0.90, P = 0.002), and pennaceous feather count (F; R = 0.33, P = 0.05) each indicating significant repeatability for females. However, none of the characteristics of the stick platform showed significant repeatability (P > 0.08).

**Table 2:** Listed nesting materials were not significantly repeatable for both female and nesting box identity. Material specified with “✝” was measured as a simple yes/no presence.

|  | Box ID |  | Female ID |  |
| --- | --- | --- | --- | --- |
|  | R & 95% CI | P | R & 95% CI | P |
| Plumulaceous | 0.03 [0 - 0.38] | 0.39 | 0.02 [0 - 0.37] | 0.43 |
| Fecal Rate | 0.07 [0 - 0.24] | < 0.99 | 0.07 [0 - 0.25] | < 0.99 |
| Spider Rate | 0.07 [0 - 0.24] | < 0.99 | 0.07 [0 - 0.23] | < 0.99 |
| † Snake Skin | <0.001 [0 - 0.88] | 0.64 | 0.14 [0 - 0.93] | 0.17 |
| † Plastic | 0.04 [0 - 0.35] | 0.32 | <0.001 [0 - 0.32] | 0.48 |

### Variation Explained By Nest Box Identity and Dimensions

Twenty-six of our boxes were used more than once over the course of the nesting season. Of the boxes, 23 were used twice while 3 were used three times. Of the boxes used more than once, the majority of the nesting attempts were from a different female than any previous attempt (22/29 total reuses; 75.9%; Table S1). With respect to general nest dimensions, the ratio of the nest height to internal box height (*R* = 0.56 [95% CI = 0.25 - 0.77], *P* < 0.001, Figure 4C), nest volume (*R =* 0.65 [95% CI = 0.37 - 0.82], *P* < 0.001), and the width of the nest’s berm (*R* = 0.49 [95% CI = 0.15 - 0.72], *P* = 0.002, Figure 4E) were all highly repeatable within a nest box. The stick platform, measured by the number of sticks in the nest, was also highly repeatable within a nest box (*R* = 0.45 [95% CI = 0.10 - 0.70], *P* = 0.008, Figure 4A).

**Figure 4:**
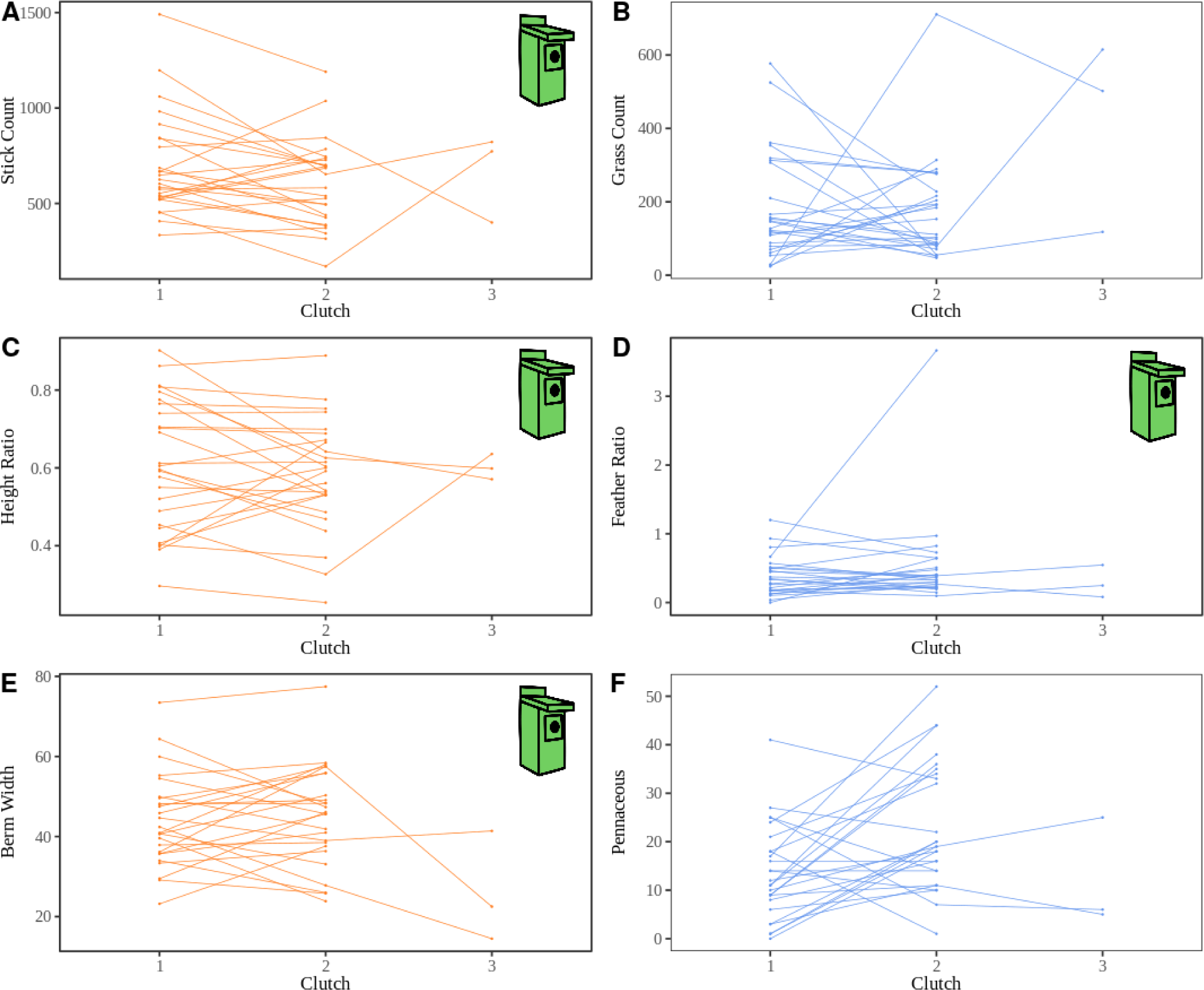
The repeatability of nest measurements across successive clutches built in the same nest box. Characteristics of the stick platform are shown in orange (left), and characteristics of the cup lining are shown in blue (right). Significant repeatability by box identity is indicated by the presence of a nest box drawing. The stick count (A; R = 0.45, P = 0.007), height ratio (C; R = 0.65, P < 0.001), berm width (E; R = 0.49, P = 0.002), and feather ratio (D; R = 0.29, P = 0.05) were significantly repeatable traits for the nesting box identity, while grass Count (B; P = 0.39) and pennaceous (F; P = 0.32) were not.

However, the components of the nest cup, the grass count (*R* = 0.04 [95% CI = 0 - 0.38], *P* = 0.38, Figure 4B), total count of feathers in the nest (*R* = 0 [95% CI = 0 - 0.34], *P* > 0.99) or the pennaceous feather count (*R* = 0.08 [95% CI = 0 - 0.42], *P* = 0.32, Figure 4F) were not statistically repeatable. The pennaceous to plumulaceous feather ratio showed moderate repeatability within a nest box (*R* = 0.29 [95% CI = 0 - 0.59], *P* = 0.05, Figure 4D), but the confidence interval overlaps with zero. The box ID did not explain the amount of spider egg sac material in the nest (*R* = 0 [95% CI = 0 - 0.21], *P* > 0.99). No other measurements were statistically repeatable (Table 2).

The height (r^2^ = 0.06, *P* = 0.006) and volume (r^2^ = 0.16, *P* < 0.001, Figure 5A) of the nest platform positively correlated with the internal height of the box. Likewise, the number of sticks within the platform positively correlated with the internal height (r^2^ = 0.12, *P* < 0.001, Figure 5B) and volume (r^2^ = 0.14, *P* < 0.001) of the box. The height of the nest was positively correlated with the height of the rectangular gap above the entrance hole (r^2^ = 0.13, *P* < 0.001, Figure 6A). It is worth noting that within our sample, boxes with taller internal heights tended to have higher rectangular gaps (r^2^ = 0.13, *P* < 0.001, Figure 6B). The width of the berm positively correlated with the diameter of the entrance hole (r² = 0.03, *P* = 0.05, Figure 7A), but not the depth of the entrance hole (*P* = 0.43, Figure 7B). The closer the top of the nest was built to the entrance hole (See Figure 1), the larger the width of the berm (r^2^ = 0.09, *P* = 0.001). No other measurement correlated with box dimensions (Table 3).

**Figure 5:**
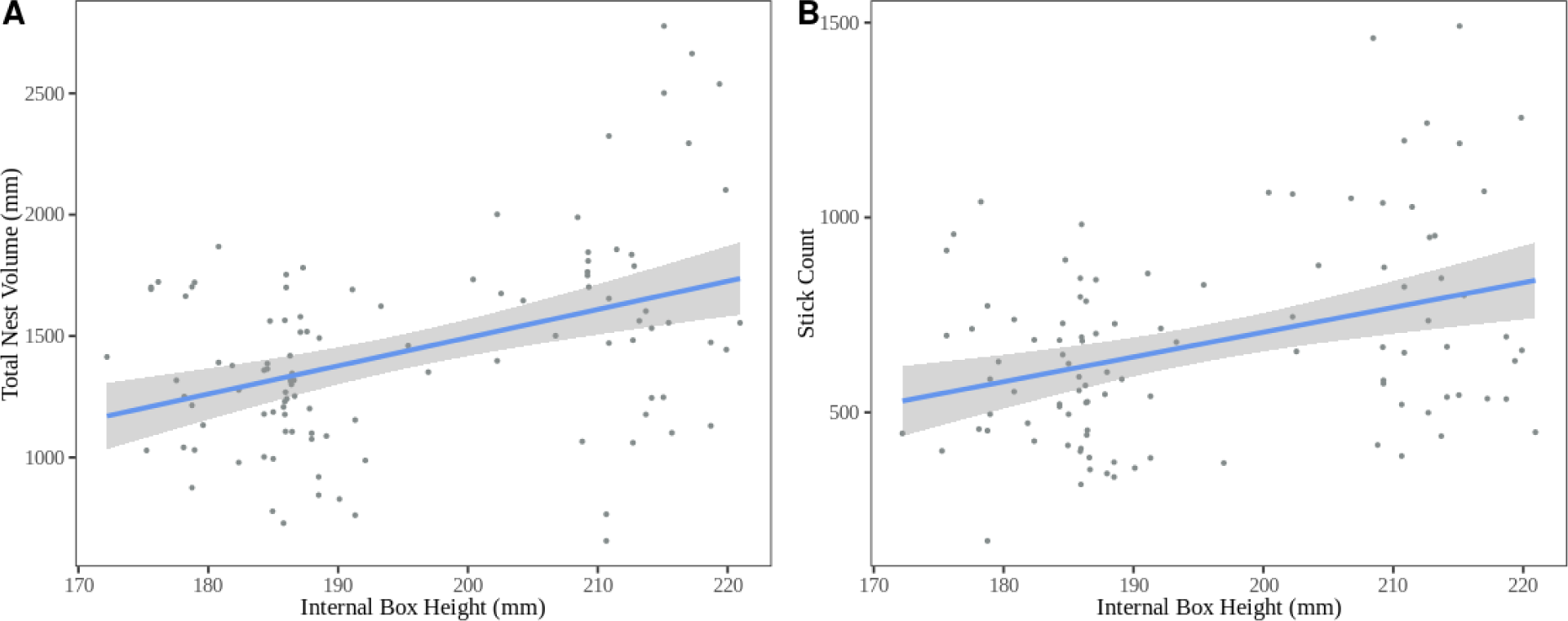
Significant positive correlations between the internal height of the nesting box and the total volume of the nest (A; r^2^ = 0.16, P < 0.001) and the stick count (B; r^2^ = 0.12, P < 0.001). Gray shading encompasses the 95% confidence interval for the line of best fit.

**Figure 6:**
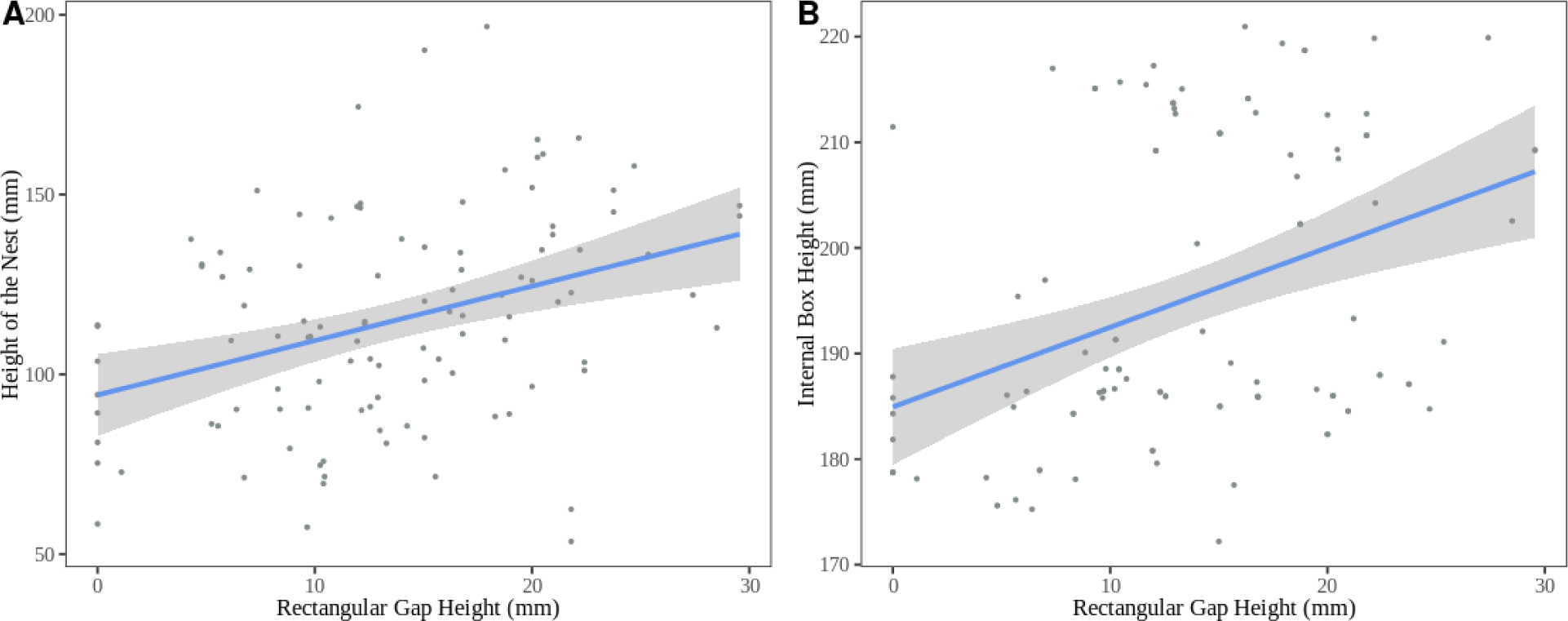
Significant positive correlations between rectangular gap height and the height of the nest (A; r^2^ = 0.13, P < 0.001) and internal box height (B; r^2^ = 0.13, P < 0.001). Gray shading encompasses the 95% confidence interval for the line of best fit.

**Figure 7:**
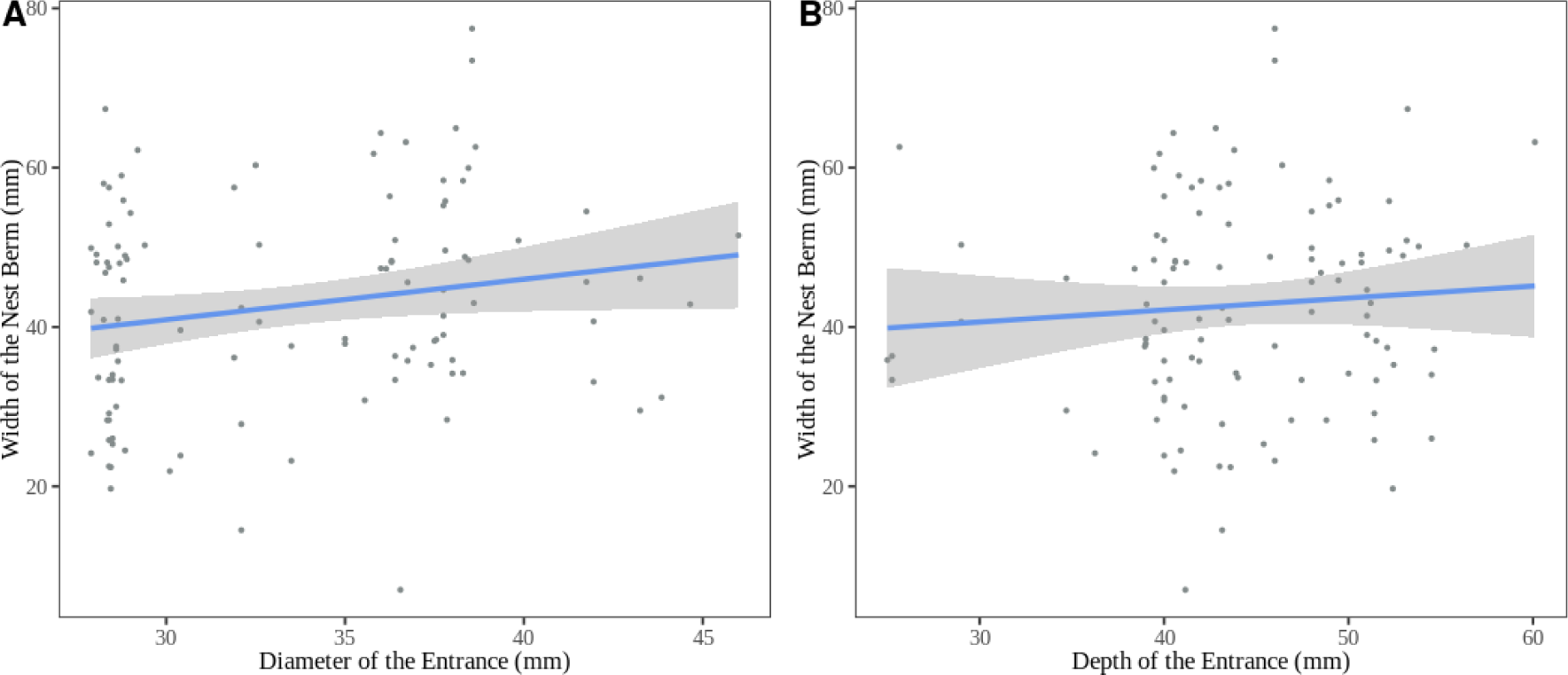
Relationships between the width of the nest berm and the diameter or depth of the entrance hole. There was a significant positive correlation between the diameter of the entrance hole and the width of the nest berm (A; r^2^ = 0.03, P = 0.05), but no detectable correlation between the depth of the entrance hole and the width of the nest berm (B; r^2^ = 0.09, P = 0.43). Gray shading encompasses the 95% confidence interval for the line of best fit.

**Table 3:**
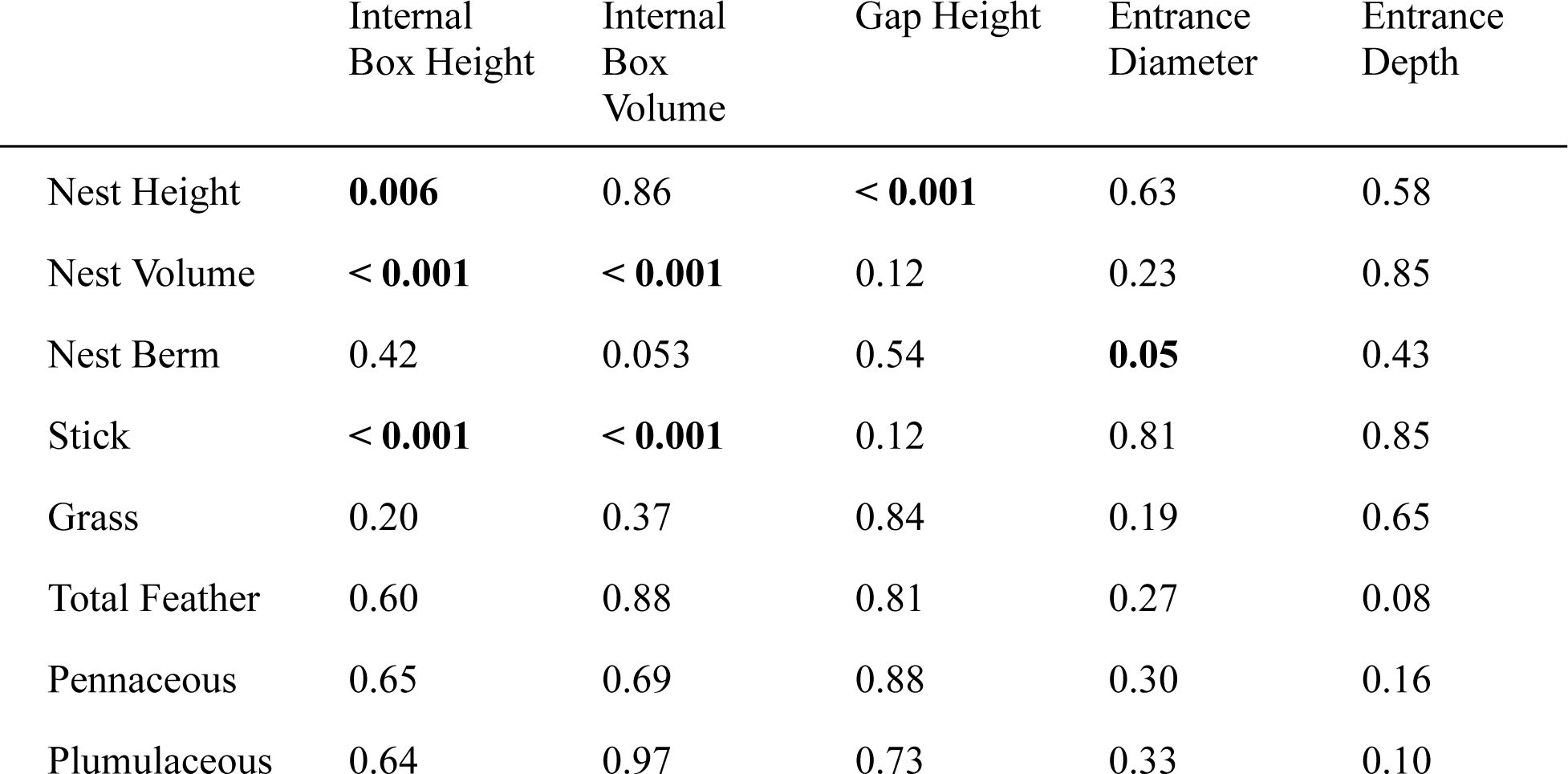
P-values of the correlations between nest characteristics and box dimensions. Significant correlations are indicated by bold text.

|  | Internal<br>Box Height | Internal<br>Box<br>Volume | Gap Height | Entrance<br>Diameter | Entrance<br>Depth |
| --- | --- | --- | --- | --- | --- |
| Nest Height | **0.006** | 0.86 | **< 0.001** | 0.63 | 0.58 |
| Nest Volume | **< 0.001** | **< 0.001** | 0.12 | 0.23 | 0.85 |
| Nest Berm | 0.42 | 0.053 | 0.54 | **0.05** | 0.43 |
| Stick | **< 0.001** | **< 0.001** | 0.12 | 0.81 | 0.85 |
| Grass | 0.20 | 0.37 | 0.84 | 0.19 | 0.65 |
| Total Feather | 0.60 | 0.88 | 0.81 | 0.27 | 0.08 |
| Pennaceous | 0.65 | 0.69 | 0.88 | 0.30 | 0.16 |
| Plumulaceous | 0.64 | 0.97 | 0.73 | 0.33 | 0.10 |

### Effects of Measurement Date and Nest Age

The date of measurement did not significantly correlate with the ratio of the nest height to internal box height (*P* = 0.54) or the width of the berm (*P* = 0.74). Neither the stick count (*P* = 0.72) or grass count (*P* = 0.88) correlated with the measurement date. A statistical trend was found with the pennaceous feather count positively correlating with the measurement date (r^2^ = 0.02, *P* = 0.07, Figure S3C), with the result reaching statistical significance with the removal of an extreme outlier (Date of year of outlier: 144; r^2^ = 0.09, *P* = 0.001), but this trend was not found with plumulaceous feathers (*P* = 0.27) or total feather count (*P* = 0.15). The amount of spider egg sacs within the nest positively correlated with the measurement date (t = 4.49, *P* < 0.001, Figure S3D). No other nest component significantly correlated with the date of measurement (*P* > 0.08, Table S3)

The pennaceous feather count positively correlated with the age of the nest (r^2^ = 0.04, *P* = 0.02, Figure S4A), but the plumulaceous (*P* = 0.43) and total feather (*P* = 0.18) counts did not. The grass count (r^2^ = 0.03, *P* = 0.054, Figure S4D) and stick count (r^2^ = 0.03, *P* = 0.052, Figure S4C) marginally negatively correlated with the age of the nest. As expected, younger nests were also less likely to contain fecal material, with nests older than 20 days almost always containing some amount of fecal material (t = 5.24, *P* < 0.001, Figure S4B). No other nest component significantly correlated with nest age (P > 0.052, Table S2).

### Reproductive Success

No correlations between clutch size or offspring success and nest morphology were observed (*P >* 0.06, Table S3). Nests sampled later in the breeding season were more likely to produce successful clutches than nests built earlier in the season (t = 2.04, *P* = 0.04, Figure S3A) but late season nests also fledged fewer offspring (t = −5.89, *P* < 0.001, Figure S3B). However, this result is likely due to females laying fewer eggs later in the season (t = −10.26, *P* < 0.001).

## DISCUSSION

### Extended Phenotypes are Complex

The niche construction hypothesis predicts that the selective pressures on construction behaviors within a species should be relatively invariable, both spatially and temporally, because the benefits of the constructed product, such as a nest, should be long term and stable (Clark et al. 2020). This invariable selection should, in turn, create consistency in the expression of the constructed product. However, we found Northern House Wren nests to be highly variable both among individuals and across a single breeding season. Consistent with previous research (Alworth 1996, Eckerle & Thompson 2005), the variation was unrelated to female body condition or the number of offspring fledged, two proxies of the builder’s fitness, suggesting that nest architecture is not under strong selection. We asked: in the absence of strong selection for specific nest characteristics, what explains the observed variability?

Extended phenotypes are a complex bidirectional interaction between the builder and their environment, with builders attempting to express their phenotype within the confines of the available space and the limitations of available building materials. We found evidence of repeatability for the female builder in multiple components of the cup composition. We also found that nest morphology was affected by the dimensions of the nesting cavity, supported by repeatability in components of successive nests built in the same box, often by different females. Our findings suggest a largely unexplored explanation for variation in nest architecture: consistent differences between individual females in their building preferences for both the types and quantity of building materials. Together with the ability of the cavity to constrain or enhance the nest dimensions, these two factors help shape the wren’s extended phenotype. We also employed an uncommon method for measuring construction behavior (i.e., directly measuring individual building materials) leading to the discovery of repeatable characteristics that would have otherwise remained hidden with more coarse measurements like nest weight. Future research should compare nest variation across populations and breeding seasons to fully understand the spatial and temporal variation of nest architecture in this species.

### Preferences in Construction behavior

Our prediction that the nesting female would be the best predictor of variation in nest architecture was only partially supported. The cup composition (i.e., the grass count, pennaceous feather count, and ratio of pennaceous to plumulaceous feathers) was moderately to highly repeatable for the female, but female identity did not explain variation in the rest of the nest morphology. This finding reinforces previous conclusions that the female constructs the nest cup independent of the male (McCabe 1965), but implicates other factors in the construction of the stick platform, such as physical constraints of the cavity. The cup appears largely unaffected by these constraints and similarly unrelated to fitness. We hypothesize that when constraints and selective pressures favoring a specific nest architecture are reduced, the effect of diverse female building preferences on nest variation is enhanced. Female Blue Tit (*Cyanistes caeruleus*) nests, for example, are also highly variable in height and this behavior is moderately repeatable for females (*R* = 0.40; Järvinen et al. 2017a) despite nest height in this species appearing unrelated to nestling survival or female body condition (Järvinen et al. 2017b). Therefore, variation between individuals in their construction preferences is likely a driving force behind much of the observed variability in nest architecture for cavity nesters.

If much of the variation in nest architecture is explained by diversity in individual preferences, it is important to understand what causes builders to differ from each other. Although this study is one of the first to tie the repeatability of an extended phenotype directly to individuals in a free-living population, lab experiments have implicated differences in experience as a potential factor that affects individual preferences. For example, Zebra Finch (*Taeniopygia castanotis*) pairs that had previous experience constructing nests built more repeatable nests during experimental trials (Whittaker et al. 2023) and juvenile black widow spiders (*Latrodectus mactans*) exposed to conspecific webs built more repeatable webs in adulthood (DiRienzo et al. 2019). Overall, having prior experiences with construction behaviors tends to result in individuals building more repeatable structures, with experiences being either previous attempts or social interactions.

Some females in our study were more consistent builders than others, but we find it unlikely that experience from prior breeding seasons could have contributed to these differences. Long-term demographic data from multiple Northern House Wren populations indicate that the majority of individuals only breed in a single season (average lifespan is 1.5 years; interannual return rates ∼30%; Johnson 2020) and so are unlikely to provide a sufficient sample size for an analysis of interannual individual differences. The age of individuals could act as a proxy of experience in some species, but we are unable to present an analysis of its effect on repeatability in this study. While molt patterns allow many songbirds to be aged as either in their first breeding season or older, this process is challenging in House Wrens (Pyle 1997), and we are not confident in our age assignments.

Individuals are also affected by trade-offs between the expression of the extended phenotype and other behaviors important for fitness. Starved colonies of *Stegodyphus sarasinorum*, for example, invest more in web construction than fed spiders (Ellendula et al. 2021), and immune stressed three-spined stickleback (*Gasterosteus aculeatus*) males construct less compact nests than healthy conspecifics (Barber et al. 2001). Thus, construction behaviors can be important indicators of the builder’s condition. We calculated the linear residual between body mass and tarsus length as a measure of female condition, but it did not correlate with the observed variability in nest cup composition. However, we collected this singular measure after the completion of egg laying, and it may not have been reflective of the female’s condition during nest building, which mostly occurs before egg laying begins (i.e., 5-7 days before our measurement). Although not measured in this study, it is likely that female body condition declined over the course of the breeding season due to the energetic costs of reproduction (Merilä and Wiggins 1997), which could also have affected nest construction. The clutch size of female House Wrens declines throughout the season, so we predicted that pairs would similarly put less effort into nest construction as the season progressed. However, we found no detectable relationships between breeding date and nest architecture. Instead, female wrens remained consistent in multiple components of the nest cup throughout the breeding season despite the fact that clutch size declines over time. Nest cup composition also did not differ between successful or unsuccessful nests or correlate with the number of eggs laid, and remained variable throughout the breeding season. Conversely, we found that nests collected longer after the initiation of construction tended to contain more pennaceous feathers than nests collected sooner, and even observed one female returning to her complete clutch of eggs with a feather in her bill, indicating a potential upkeep behavior in cup composition despite the increasing needs of the offspring. The lack of any apparent reproductive cost of the nest architecture conflicts with the results of other studies on extended phenotypes. For example, Zebra Finches modify successive nests when the previous clutch fails (Edwards et al. 2020). However, this finding remains consistent with previous literature on Northern House Wrens (Alworth 1996, Pribil & Picman 1997, Eckerle & Thompson 2005). We discuss potential causes for this lack of fitness-related nest architecture below.

### Physical Constraints and Environmental Pressures on Constructions

The box cavity was also a significant predictor of nest morphology and composition, specifically that of the stick platform. This repeatability supports the prediction that nest architecture would be related to constraints posed by the cavity dimensions. House Wrens appeared to be filling the available space and adjusting nest morphology based on differences between box openings. Stanback et al. (2013) suggested that wrens were modulating the nest structure to accommodate for perceived vulnerabilities in the cavity, and our data support this conclusion. For example, the size of the entrance hole is a potential source of vulnerability, as more accessible nests are more likely to be depredated and parasitized (Paclík et al. 2009, Hauber et al. 2024). Alternatively, larger entrances could increase heat loss, increasing the threat of hypothermia for the offspring (Lamprecht and Schmolz 2004). We found that House Wrens built wider berms in boxes with larger entrance hole diameters, potentially modulating their nest structure to better protect the contents of the nest from predators or to increase heat retention. We found a similar correlation between the height of the rectangular gap and the height of the nest, as larger gaps resulted in taller nests. Previous research has suggested that the berm could act as a barrier between the nestlings within the cup and any outside threats (Stanback et al. 2013), however evidence for this relationship is inconsistent (Pribil & Picman, 1997). Despite our support of Stanback et al.’s (2013) finding, the width of the nest berm had no effect on the number of eggs laid or nestling survival. Variation in the width of the berm also did not change throughout the breeding season, despite the prediction that nest architecture that reduces insulation would be favored as temperatures increased. House Wrens may be modulating their construction behaviors in response to perceived vulnerability, but our results suggest that this action has little to no effect on nest success. Previous literature has implicated the climate in affecting the evolution of nest characteristics, but this relationship appears to be affecting nest shape at the among- rather than within-species level (e.g., domed v. open-cup nesting species) and still explains only a small portion of the observed variation (Colombo et al. 2024).

However, evidence from other species still suggests that structures protect inhabitants from predators and environmental exposure by the construction of a physical barrier. For example, eurasian beavers (*Castor fiber*) in the Netherlands, which face lower predation rates and milder climates than their northern counterparts, build lodges with thinner walls (Rosell & Campbell-Palmer 2022). Cavity-nesters are unique in that they do not need to construct a physical barrier, as the cavity itself is the barrier between the inhabitants and the outside environment. Therefore, we hypothesize that the lack of any detectable relationship between nest architecture and fitness in House Wrens could be due to selection favoring a better quality cavity rather than specific characteristics of the nest. Observations of platypus (*Ornithorhynchus anatinus*) nests, which are housed inside burrows, are consistent with this hypothesis as the dimensions of the burrow explained offspring production, while effort in nest construction was unrelated (Thomas et al. 2018). Support for this hypothesis in House Wrens is more indirect, but males with empty cavities due to experimentally removed nests were just as likely to acquire a mate as control males that initiated nest building (Alworth 1996), suggesting that the cavity rather than the contents are more important in mate choice. Female House Wrens also use cavity quantity before male quality when choosing mates (Eckerle & Thompson, 2006) and male House Wrens selectively choose nesting boxes with smaller entrances that likely provide superior protection from predators and brood parasites (Pribil & Picman 1997). Collectively, the available data suggest that the reproductive success of House Wrens is more related to the cavity itself than the nest.

### Limitations

Extended phenotypes become additionally complex when more than one individual contributes to the construction effort. Both sexes contribute to the nest building in House Wrens, so it is important to consider how the male contribution may affect our data and interpretation. Although their contributions are likely not equal, with the female completing the majority of the nest (Alworth & Scheiber 2000), we were unable to directly account for the effect of individual males on construction behavior because the majority of males were unbanded. Despite this limitation, Northern House Wren pairs often change between nesting attempts (Baldwin 1921, Drilling & Thompson 1988), so it is rarely the same pair building successive nests. In 2019, we found that out of 31 banded pairs in our population, only 6 (19.4%) remained with the same mate for the entire breeding season, with the female being more likely to leave a territory than the male. The fact that females were repeatable in cup composition, despite frequent mate switching, supports our conclusion that the female constructs some aspects of the nest independently of the male. Box identity could potentially be a proxy of the male identity, as cavities are controlled by individual males on their own territories; however, males often control many boxes within the same territory and the rate of movement between these boxes is unknown. Therefore, we are unable to split apart the identity of the male from the identity of the box in this study. Males begin bringing sticks to the cavity before acquiring a mate (Johnson 2020), and so if the box identity is a reliable proxy for male identity, then the male contribution to building could help explain the observed variation in the stick platform, but more research is needed.

In most cases, we only measured the construction attempts of individuals twice. House Wrens only produce two to three clutches a season, so we are limited by this number and the likelihood the female renested within our nest boxes instead of natural cavities. However, Bell et al. (2009) found no significant effect of the number of observations per individual on repeatability estimates. Instead, the number of observations appears to have a greater effect on the range of uncertainty in the estimate. We provided this range of uncertainty, as a confidence interval, and required the lower end to remain higher than zero for significance. Furthermore, all of our estimates are calculated from a sample size (*N* = 24 for female identity; *N* = 26 for box identity) larger than the median sample size (*N* = 22) of all studies examined by Bell et al. (2009), and so we do not believe that our sample size is a major limitation.

### Conclusions

The repeatability of construction behaviors is greatly understudied, particularly in free-living populations. We examined the Northern House Wren, a species with incredible variation in nest architecture (See Figure S1), to estimate the amount of variability in construction behavior explained by the identity of the female builder and the box identity and its dimensions. We found female northern House Wrens to be moderately to highly repeatable in the composition of the nest cup, but this behavior was unrelated to female body condition or nestling survival. The general dimensions of the nest, as well as the stick platform, correlated with the dimensions of the box, and these measurements were highly repeatable for box identity. These results support previous findings that House Wrens modify aspects of the nest to compensate for vulnerabilities in the cavity. We hypothesize that the observed variability in nest architecture is due to the nest architecture being under minimal selective pressure, thus allowing the diversity of female building preferences to have a greater effect on the expression of the phenotype. Instead, choosing a protective cavity is likely to be more important for the builder’s fitness, with the nest construction process being relatively unimportant for mate selection or nestling survival. However, much of House Wren nest construction still remains unclear, including the contribution of the male in constructing the stick platform. Extended phenotypes are complex, requiring an examination of both the environment *and* the builder. The cause of variation in House Wren nest morphology has been largely unexplained due to a narrow focus on potential external factors. Studying the repeatability of extended phenotypes across taxa has provided new insights into the construction behaviors of animals and provides an important method for understanding the causes and consequences of individual preferences.

## Data Availability Statement

The data and code that support these findings are available in the Mendeley Data Repository at doi: 10.17632/fnhr5mm99p.1.

## ACKNOWLEDGMENTS

We thank Ohio Wesleyan University, the Big Walnut School System, and the Davis, Koban, and Fink families for access to facilities and field sites. We also like to thank Ali Amer for helping in nest dissection, Ava Swanson for the creation of the House Wren and box drawings used in Figures 3 and 7, and Martin Stoffel for statistical advice. Finally, we would like to thank the House Wrens for making our work possible. This project was funded by the Ohio Wesleyan University Summer Science Research Program.

All procedures were reviewed and approved by the Ohio Wesleyan University Institutional Animal Care and Use Committee (Protocol Number: 05-2021-06), U.S. Federal Bird Banding Lab (Permit Number: 24035), and State of Ohio Department of Natural Resources (Permit Numbers: 23-009, 23-010).

## SUPPLEMENTARY

**Figure S1:**
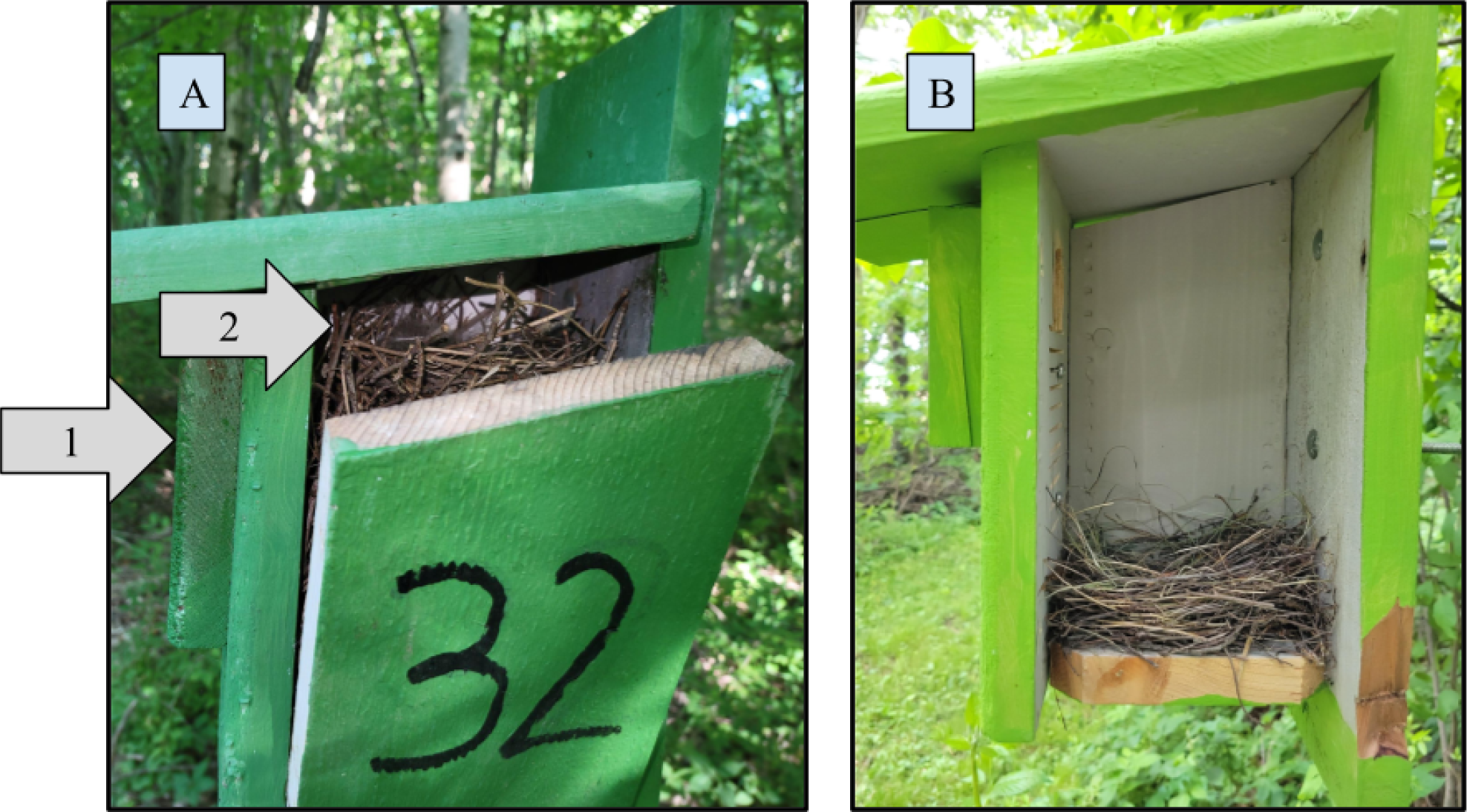
Images of complete nests constructed in a nesting box in 2022 showing that A) nests are sometimes constructed to the point where they will cover the primary entrance of the nesting box, while B) some nests are constructed only a few centimeters tall. Arrow 1 points to the top of the primary entrance; Arrow 2 points to the top of the nest. In cases like that shown in nest A, the wrens will use the gap entrance (See Figure 1) as their main entrance instead.

**Figure S2:**
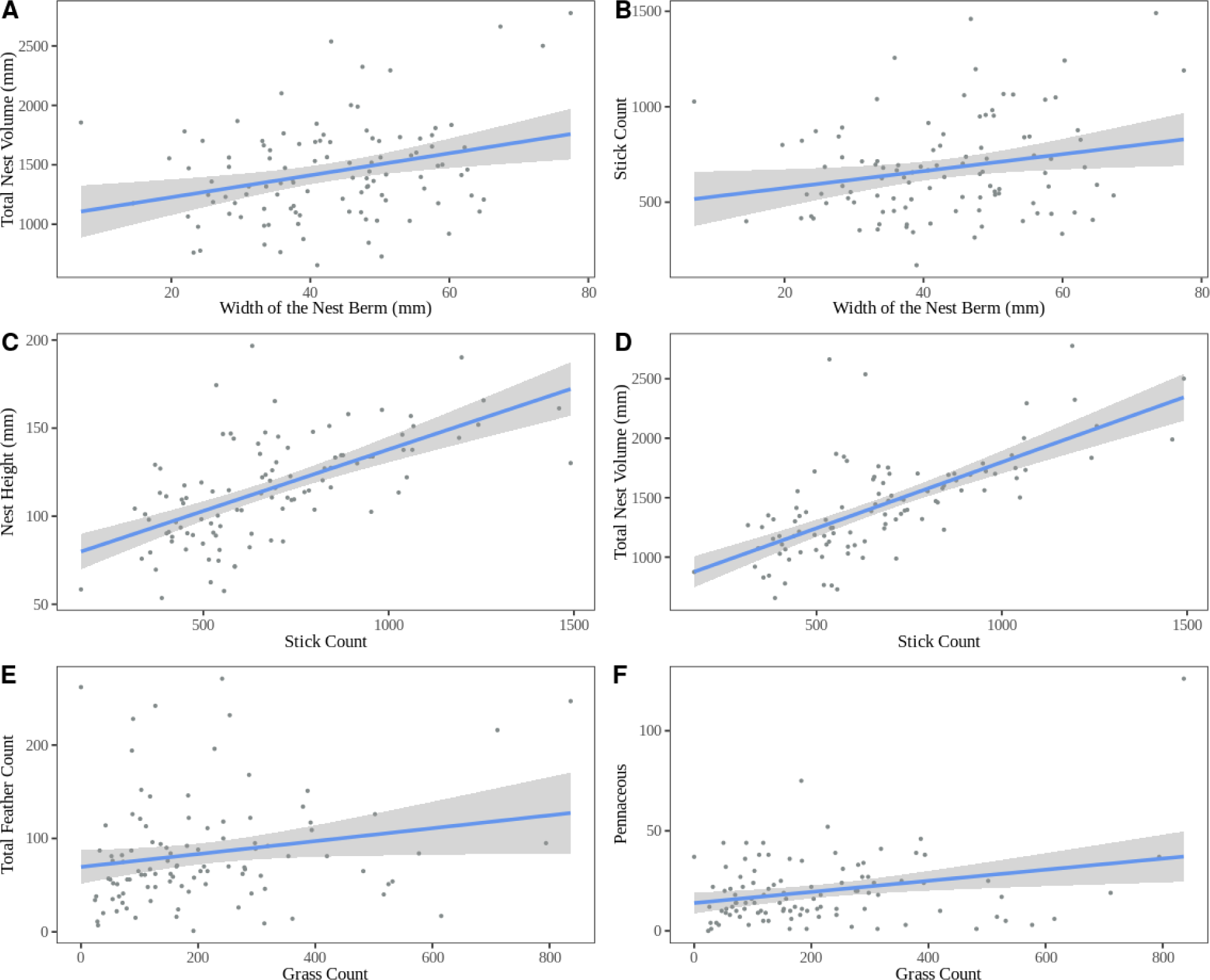
Relationships between different nest attributes and building materials. The width of the nest berm was positively correlated with the total volume (A; r^2^ = 0.09, P < 0.001) and stick count (B; r^2^ = 0.04, P = 0.02) of the nest. The stick count further highly positively correlated with the height of the nest (C; r^2^ = 0.37, P < 0.001) and the total volume of the nest (D; r^2^ = 0.41, P < 0.001). The grass count positively correlated with the total count of feathers (E; r^2^ = 0.03, P = 0.04) and pennaceous feather count (F; r^2^ = 0.06, P = 0.005). Gray shading encompasses the 95% confidence interval for the line of best fit.

**Figure S3:**
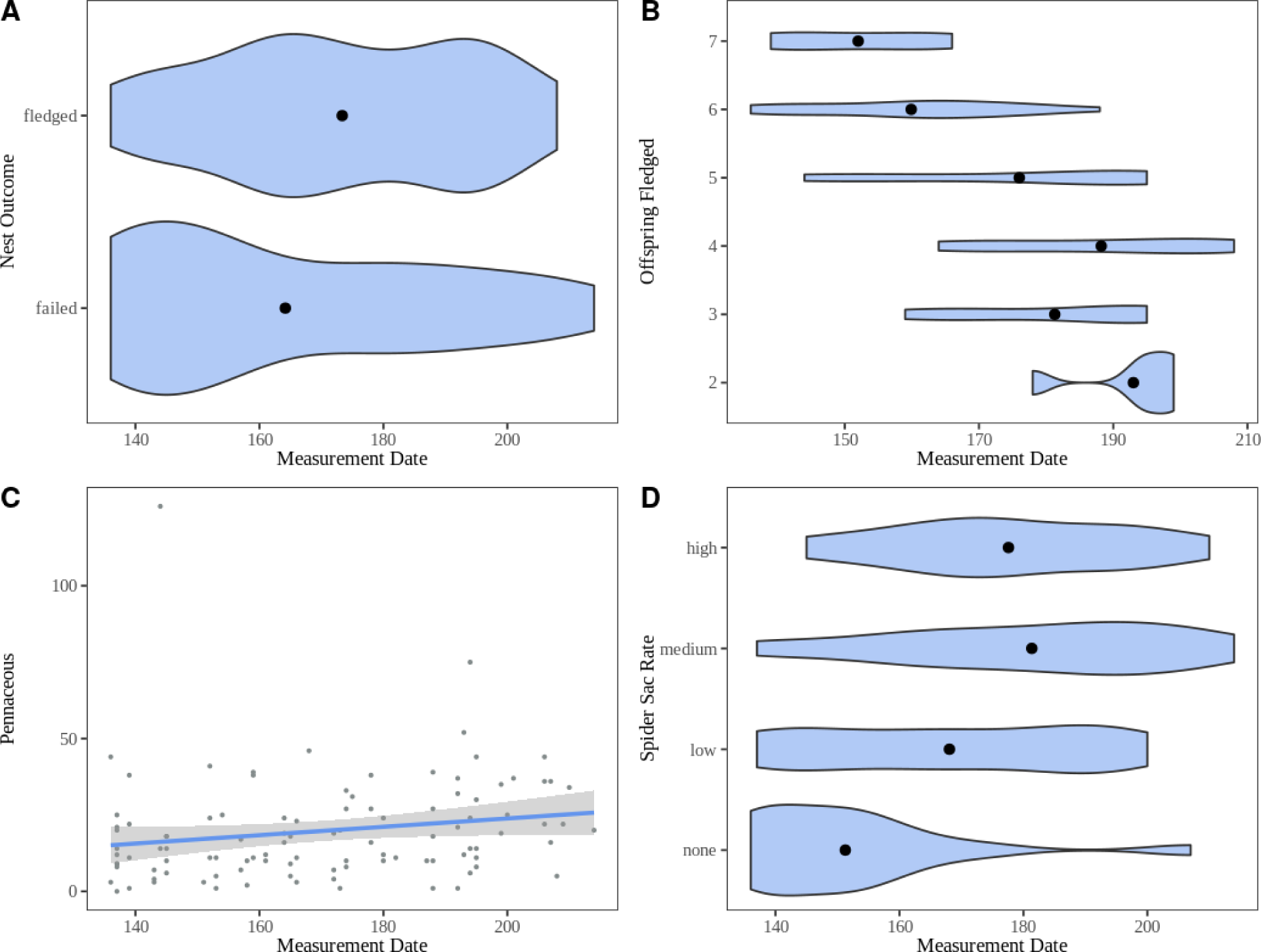
Significant relationships between nest characteristics and the date of nest measurement (i.e., completion of egg laying). Nests measured earlier in the season were more likely to fail than nests measured later in the season (A; t = 2.04, P = 0.04), but also fledged fewer offspring (B; t = −5.89, P = 0.002). The pennaceous feather count positively correlated with the measurement date after the removal of the outlier on DOY 144 (C; r^2^ = 0.02, P = 0.001). Nests measured later in the season also contained more spider sac material than those measured earlier in the season (D; t = 4.49, P < 0.001). Gray shading in C encompasses the 95% confidence interval for the line of best fit.

**Figure S4:**
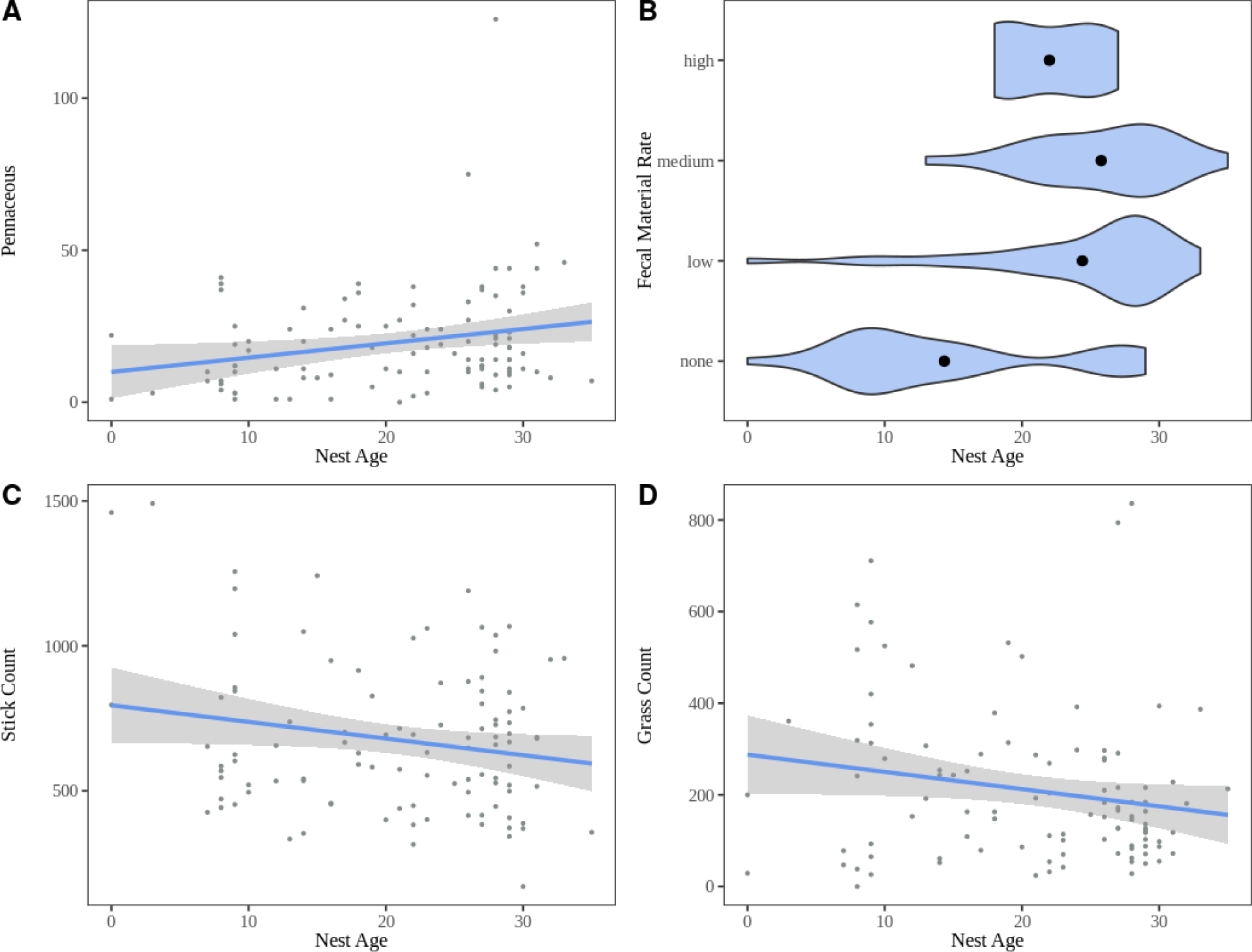
Relationship between three nest materials and fecal material with the age of the nest at collection. Pennaceous feather count positively correlated with the nest age (A; r^2^ = 0.04, P = 0.02). Although not statistically significant, the stick count (C; r^2^ = 0.03, P = 0.052) and the grass count (D; r^2^ = 0.03, P = 0.054) negatively correlated with the age of the nest. Older nests contained more fecal material than younger nests (B; t = 5.24, P < 0.001). Gray shading in A, C, and D encompasses the 95% confidence interval for the line of best fit.

**Table S1:**
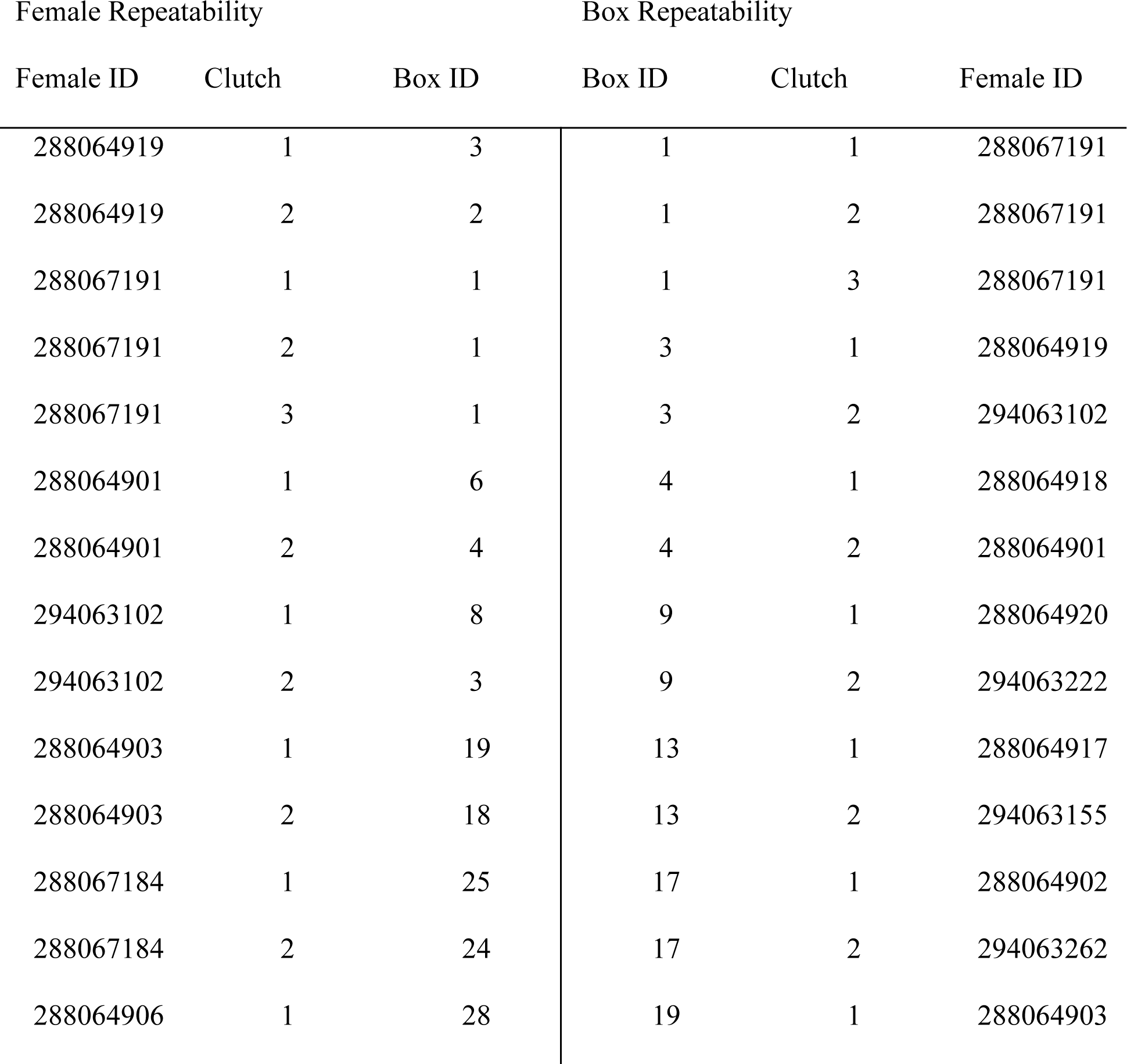
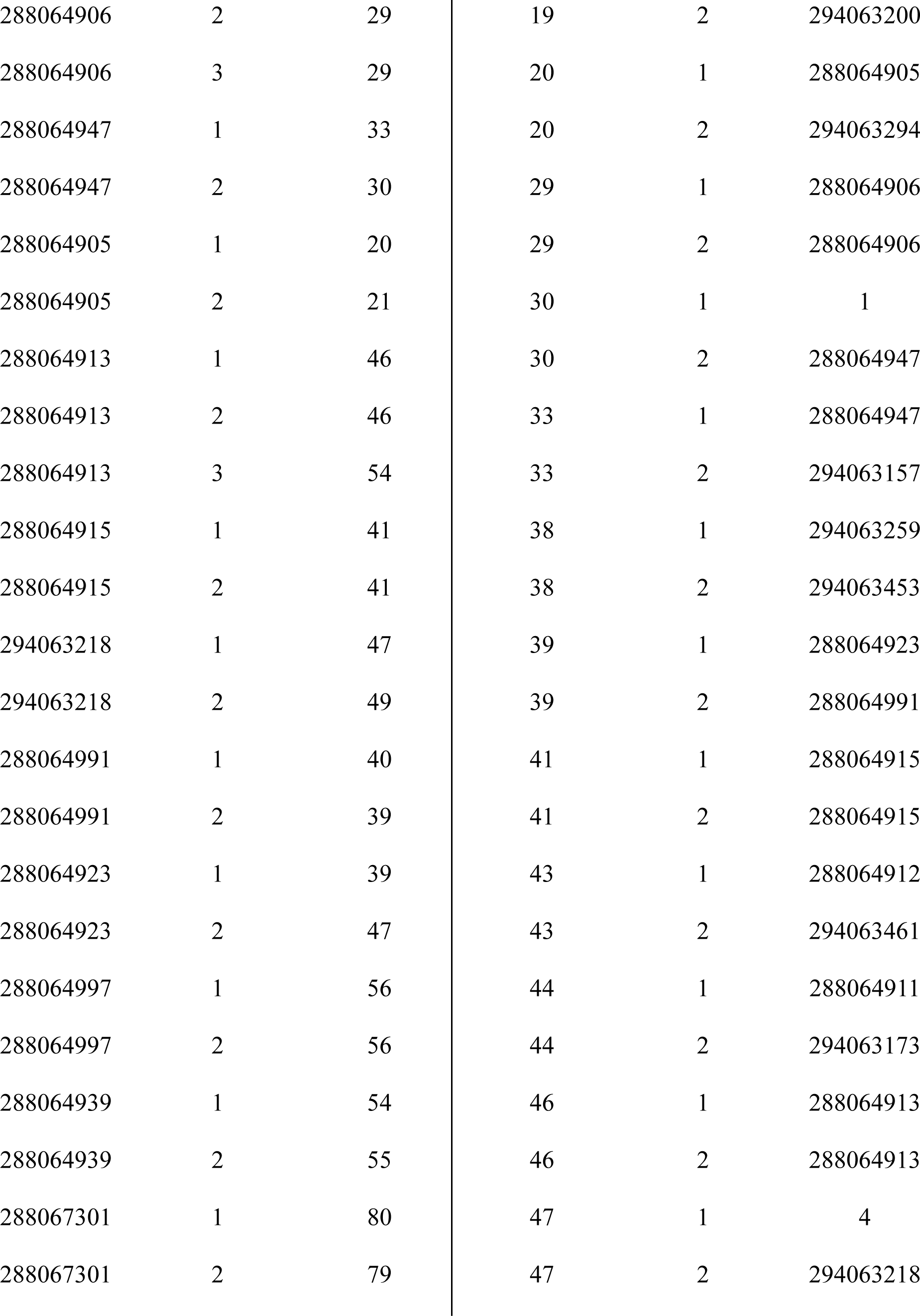
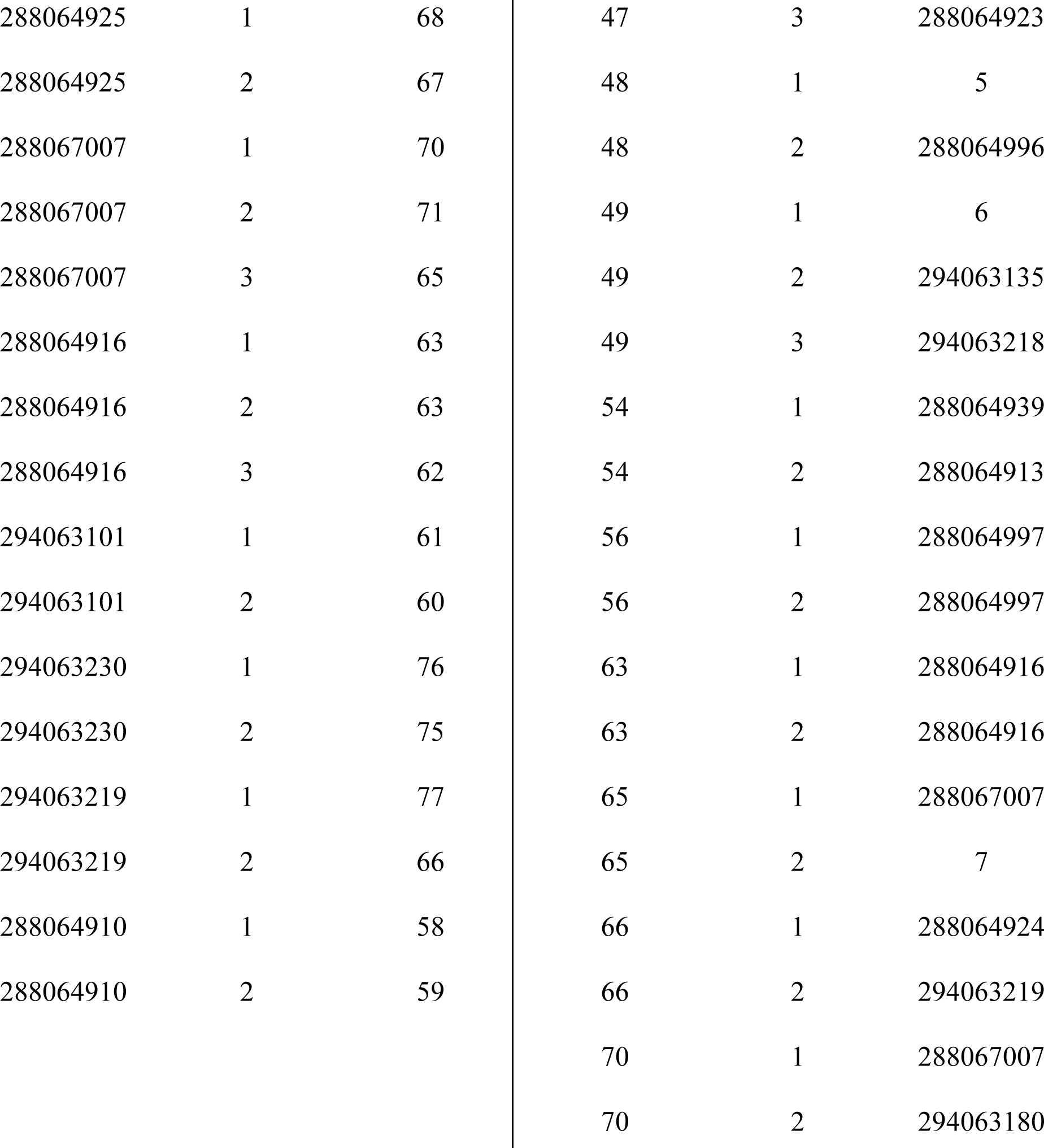
Individuals examined for Female Identity repeatable traits and Box Identity traits. Females with single IDs were unbanded individuals.

**Table S2:**
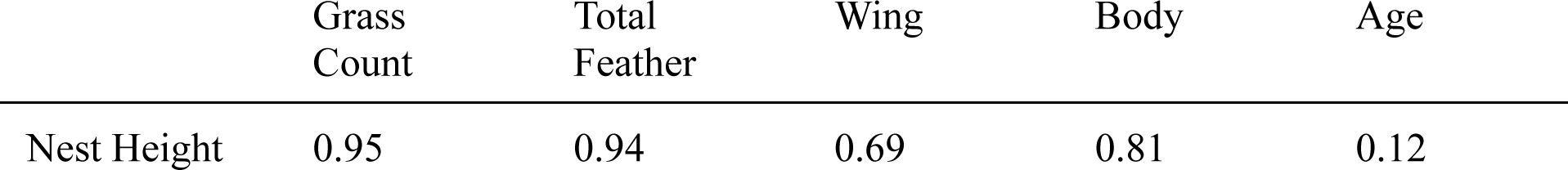
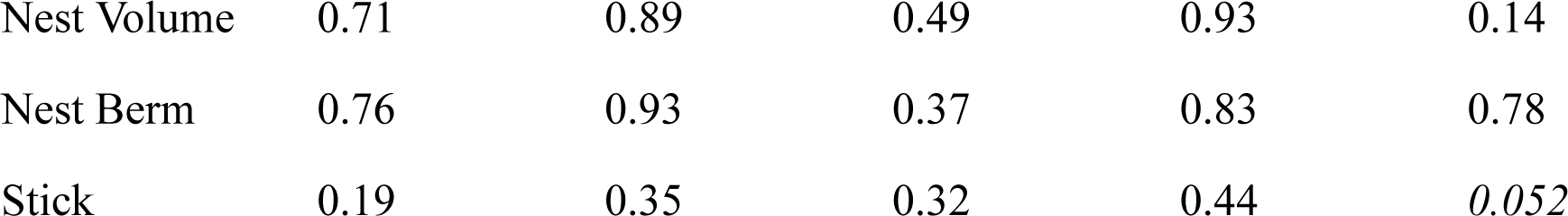
P-values supporting the lack of significant correlations between nest characteristics and the nest age.

**Table S3:**
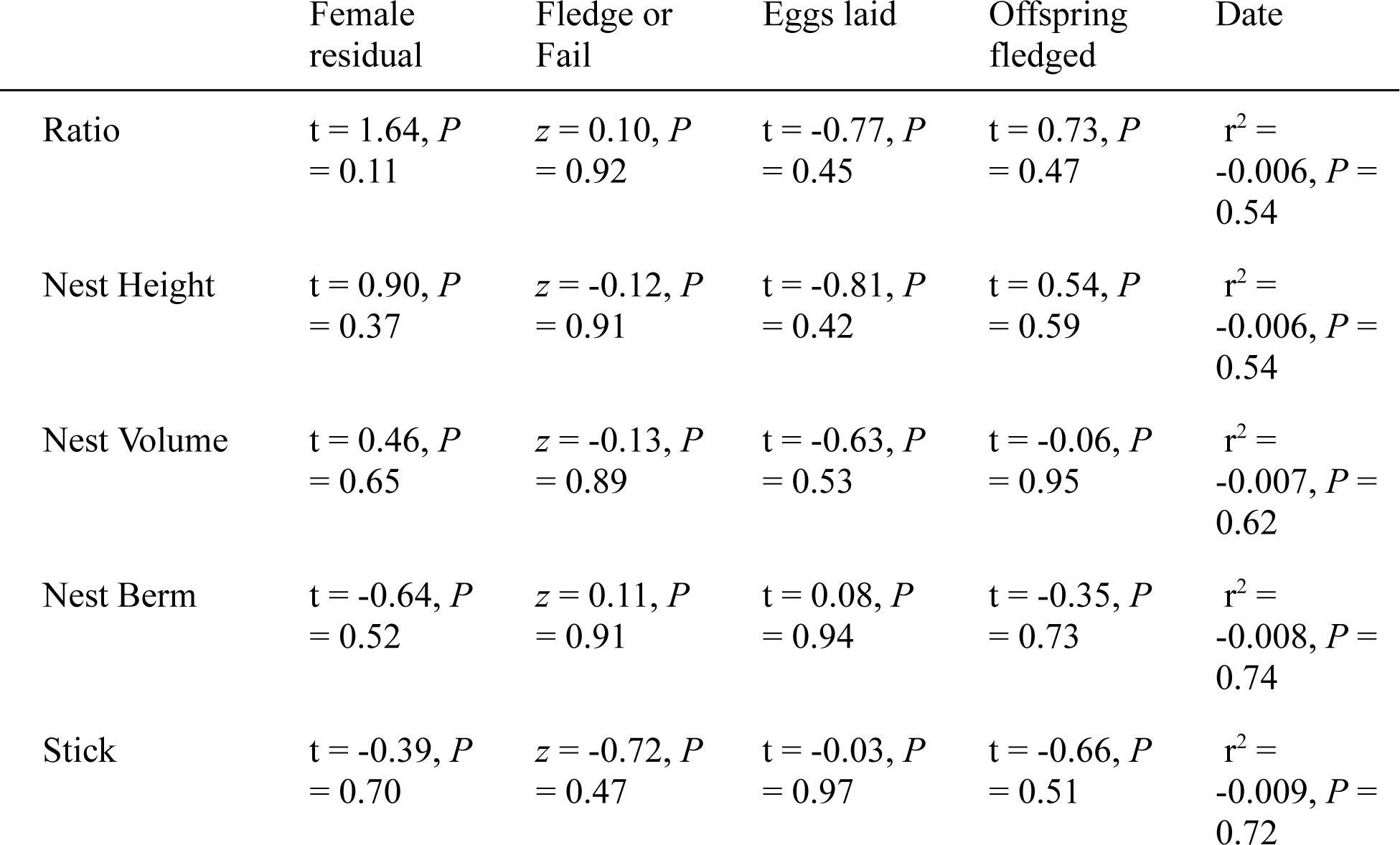
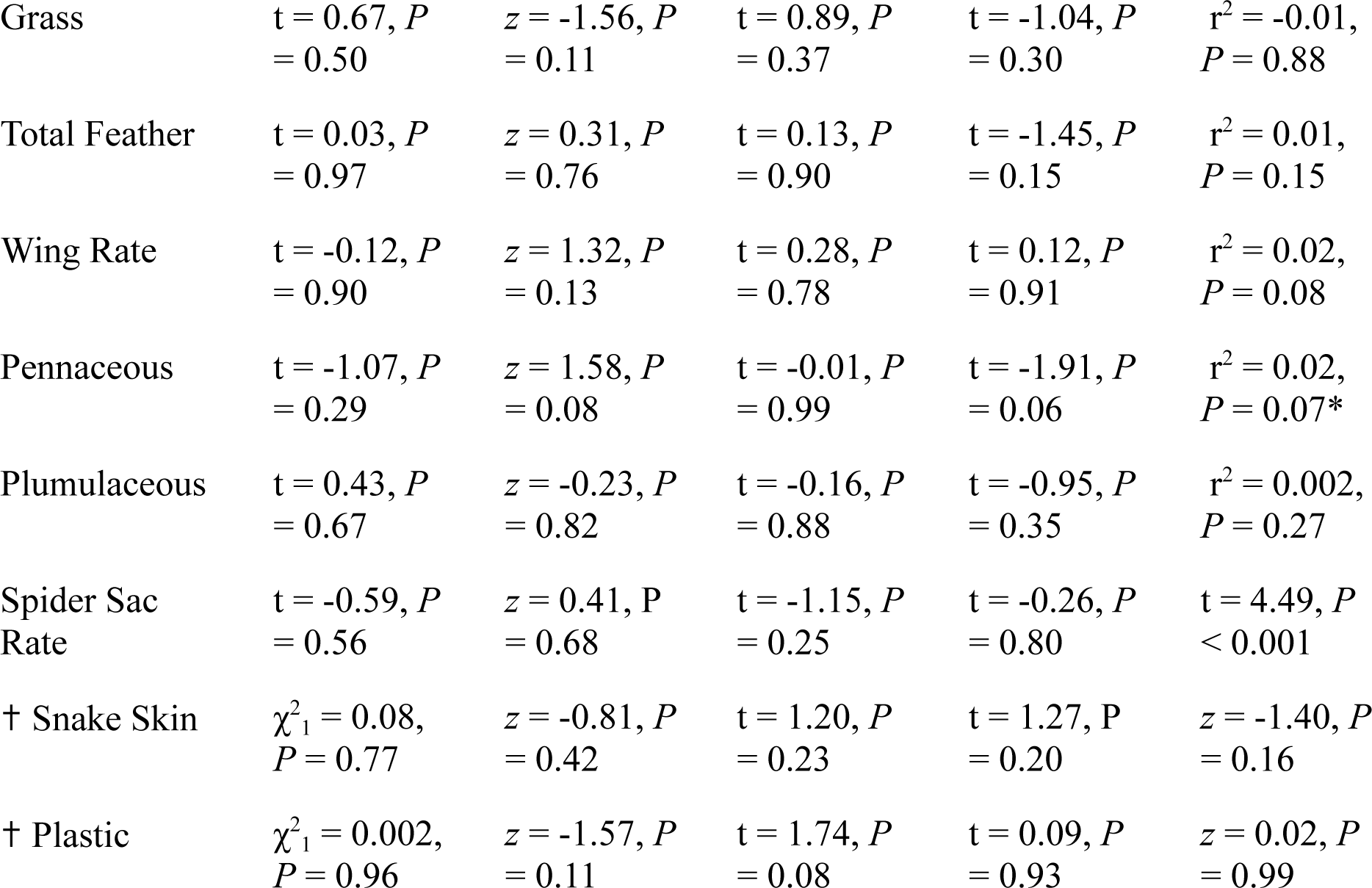
Relationships between the dimensions and components of the nest and various proxies of fitness: Female residual - defined by the correlation between the tarsus length and the body mass, Fledge or Fail - the simply outcome of the nest, Eggs laid - the number of eggs originally laid by the nesting female at the completion of the cup, and Offspring fledged - the estimated number of offspring that fledged the nest (only successful nests examined). Date - the relationship between the dimensions and components of the nest and the date at which the nest was measured. ✝Specified material measured as a simple yes/no presence. * r^2^ = 0.09, P = 0.001 with removal of outlier date 144

