## Supplementary Information for "Repeatability of an extended phenotype: potential causes and consequences of nest variation in the Northern House Wren (Troglodytes aedon aedon)"

*Table S1 - Individuals examined for Female Identity repeatable traits and Box Identity traits. Females with single IDs were unbanded individuals.*

| Female Repeatability | | | Box Repeatability | | |
|---|---|---|---|---|---|
| Female ID | Clutch | Box ID | Box ID | Clutch | Female ID |
| 288064919 | 1 | 3 | 1 | 1 | 288067191 |
| 288064919 | 2 | 2 | 1 | 2 | 288067191 |
| 288067191 | 1 | 1 | 1 | 3 | 288067191 |
| 288067191 | 2 | 1 | 3 | 1 | 288064919 |
| 288067191 | 3 | 1 | 3 | 2 | 294063102 |
| 288064901 | 1 | 6 | 4 | 1 | 288064918 |
| 288064901 | 2 | 4 | 4 | 2 | 288064901 |
| 294063102 | 1 | 8 | 9 | 1 | 288064920 |
| 294063102 | 2 | 3 | 9 | 2 | 294063222 |
| 288064903 | 1 | 19 | 13 | 1 | 288064917 |
| 288064903 | 2 | 18 | 13 | 2 | 294063155 |
| 288067184 | 1 | 25 | 17 | 1 | 288064902 |
| 288067184 | 2 | 24 | 17 | 2 | 294063262 |
| 288064906 | 1 | 28 | 19 | 1 | 288064903 |
| 288064906 | 2 | 29 | 19 | 2 | 294063200 |
| 288064906 | 3 | 29 | 20 | 1 | 288064905 |
| 288064947 | 1 | 33 | 20 | 2 | 294063294 |
| 288064947 | 2 | 30 | 29 | 1 | 288064906 |

| | | | | | |
|---|---|---|---|---|---|
| 288064905 | 1 | 20 | 29 | 2 | 288064906 |
| 288064905 | 2 | 21 | 30 | 1 | 1 |
| 288064913 | 1 | 46 | 30 | 2 | 288064947 |
| 288064913 | 2 | 46 | 33 | 1 | 288064947 |
| 288064913 | 3 | 54 | 33 | 2 | 294063157 |
| 288064915 | 1 | 41 | 38 | 1 | 294063259 |
| 288064915 | 2 | 41 | 38 | 2 | 294063453 |
| 294063218 | 1 | 47 | 39 | 1 | 288064923 |
| 294063218 | 2 | 49 | 39 | 2 | 288064991 |
| 288064991 | 1 | 40 | 41 | 1 | 288064915 |
| 288064991 | 2 | 39 | 41 | 2 | 288064915 |
| 288064923 | 1 | 39 | 43 | 1 | 288064912 |
| 288064923 | 2 | 47 | 43 | 2 | 294063461 |
| 288064997 | 1 | 56 | 44 | 1 | 288064911 |
| 288064997 | 2 | 56 | 44 | 2 | 294063173 |
| 288064939 | 1 | 54 | 46 | 1 | 288064913 |
| 288064939 | 2 | 55 | 46 | 2 | 288064913 |
| 288067301 | 1 | 80 | 47 | 1 | 4 |
| 288067301 | 2 | 79 | 47 | 2 | 294063218 |
| 288064925 | 1 | 68 | 47 | 3 | 288064923 |
| 288064925 | 2 | 67 | 48 | 1 | 5 |
| 288067007 | 1 | 70 | 48 | 2 | 288064996 |
| 288067007 | 2 | 71 | 49 | 1 | 6 |

| | | | | | |
|---|---|---|---|---|---|
| 288067007 | 3 | 65 | 49 | 2 | 294063135 |
| 288064916 | 1 | 63 | 49 | 3 | 294063218 |
| 288064916 | 2 | 63 | 54 | 1 | 288064939 |
| 288064916 | 3 | 62 | 54 | 2 | 288064913 |
| 294063101 | 1 | 61 | 56 | 1 | 288064997 |
| 294063101 | 2 | 60 | 56 | 2 | 288064997 |
| 294063230 | 1 | 76 | 63 | 1 | 288064916 |
| 294063230 | 2 | 75 | 63 | 2 | 288064916 |
| 294063219 | 1 | 77 | 65 | 1 | 288067007 |
| 294063219 | 2 | 66 | 65 | 2 | 7 |
| 288064910 | 1 | 58 | 66 | 1 | 288064924 |
| 288064910 | 2 | 59 | 66 | 2 | 294063219 |
| | | | 70 | 1 | 288067007 |
| | | | 70 | 2 | 294063180 |

*Table S2 - P-values supporting the lack of significant correlations between nest characteristics and the nest age.*

| | Grass Count | Total Feather | Wing | Body | Age |
|---|---|---|---|---|---|
| Nest Height | 0.95 | 0.94 | 0.69 | 0.81 | 0.12 |
| Nest Volume | 0.71 | 0.89 | 0.49 | 0.93 | 0.14 |
| Nest Berm | 0.76 | 0.93 | 0.37 | 0.83 | 0.78 |
| Stick | 0.19 | 0.35 | 0.32 | 0.44 | *0.052* |

*Table S3 - Relationships between the dimensions and components of the nest and various proxies of fitness: Female residual - defined by the correlation between the tarsus length and the body mass, Fledge or Fail - the simply outcome of the nest, Eggs laid - the number of eggs originally laid by the nesting female at the completion of the cup, and Offspring fledged - the estimated number of offspring that fledged the nest (only successful nests examined). Date - the relationship between the dimensions and components of the nest and the date at which the nest was measured. †Specified material measured as a simple yes/no presence. * $r^2 = 0.09$, $P = 0.001$ with removal of outlier date 144*

| | Female residual | Fledge or Fail | Eggs laid | Offspring fledged | Date |
|---|---|---|---|---|---|
| Ratio | t = 1.64, *P* = 0.11 | *z* = 0.10, *P* = 0.92 | t = -0.77, *P* = 0.45 | t = 0.73, *P* = 0.47 | $r^2$ = -0.006, *P* = 0.54 |
| Nest Height | t = 0.90, *P* = 0.37 | *z* = -0.12, *P* = 0.91 | t = -0.81, *P* = 0.42 | t = 0.54, *P* = 0.59 | $r^2$ = -0.006, *P* = 0.54 |
| Nest Volume | t = 0.46, *P* = 0.65 | *z* = -0.13, *P* = 0.89 | t = -0.63, *P* = 0.53 | t = -0.06, *P* = 0.95 | $r^2$ = -0.007, *P* = 0.62 |
| Nest Berm | t = -0.64, *P* = 0.52 | *z* = 0.11, *P* = 0.91 | t = 0.08, *P* = 0.94 | t = -0.35, *P* = 0.73 | $r^2$ = -0.008, *P* = 0.74 |
| Stick | t = -0.39, *P* = 0.70 | *z* = -0.72, *P* = 0.47 | t = -0.03, *P* = 0.97 | t = -0.66, *P* = 0.51 | $r^2$ = -0.009, *P* = 0.72 |
| Grass | t = 0.67, *P* = 0.50 | *z* = -1.56, *P* = 0.11 | t = 0.89, *P* = 0.37 | t = -1.04, *P* = 0.30 | $r^2$ = -0.01, *P* = 0.88 |
| Total Feather | t = 0.03, *P* = 0.97 | *z* = 0.31, *P* = 0.76 | t = 0.13, *P* = 0.90 | t = -1.45, *P* = 0.15 | $r^2$ = 0.01, *P* = 0.15 |
| Wing Rate | t = -0.12, *P* = 0.90 | *z* = 1.32, *P* = 0.13 | t = 0.28, *P* = 0.78 | t = 0.12, *P* = 0.91 | $r^2$ = 0.02, *P* = 0.08 |

| | | | | | |
|---|---|---|---|---|---|
| Pennaceous | t = -1.07, *P* = 0.29 | *z* = 1.58, *P* = 0.08 | t = -0.01, *P* = 0.99 | t = -1.91, *P* = 0.06 | $r^2$ = 0.02, *P* = 0.07* |
| Plumulaceous | t = 0.43, *P* = 0.67 | *z* = -0.23, *P* = 0.82 | t = -0.16, *P* = 0.88 | t = -0.95, *P* = 0.35 | $r^2$ = 0.002, *P* = 0.27 |
| Spider Sac Rate | t = -0.59, *P* = 0.56 | *z* = 0.41, P = 0.68 | t = -1.15, *P* = 0.25 | t = -0.26, *P* = 0.80 | t = 4.49, *P* < 0.001 |
| † Snake Skin | $\chi^2_1$ = 0.08, *P* = 0.77 | *z* = -0.81, *P* = 0.42 | t = 1.20, *P* = 0.23 | t = 1.27, P = 0.20 | *z* = -1.40, *P* = 0.16 |
| † Plastic | $\chi^2_1$ = 0.002, *P* = 0.96 | *z* = -1.57, *P* = 0.11 | t = 1.74, *P* = 0.08 | t = 0.09, *P* = 0.93 | *z* = 0.02, *P* = 0.99 |

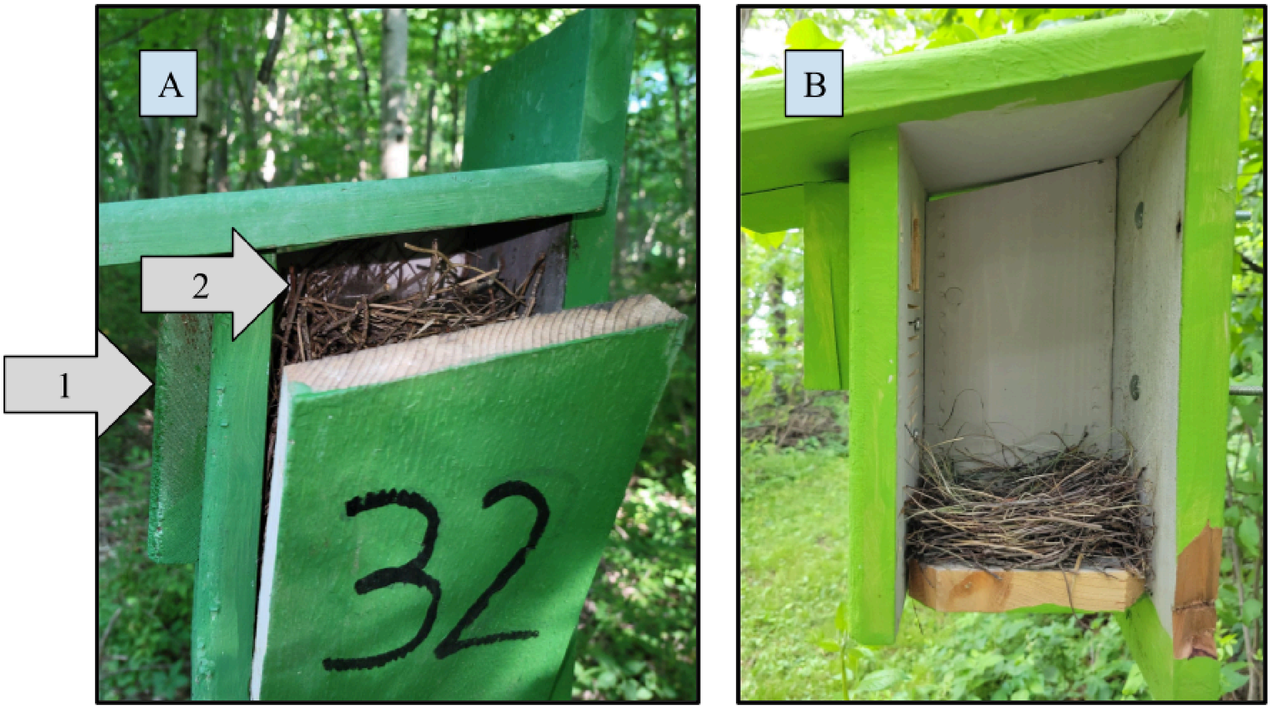


*Figure S1 - Images of complete nests constructed in a nesting box in 2022 showing that A) nests are sometimes constructed to the point where they will cover the primary entrance of the nesting box, while B) some nests are constructed only a few centimeters tall. Arrow 1 points to*

*the top of the primary entrance; Arrow 2 points to the top of the nest. In cases like that shown in nest A, the wrens will use the gap entrance (See Figure 1) as their main entrance instead.*

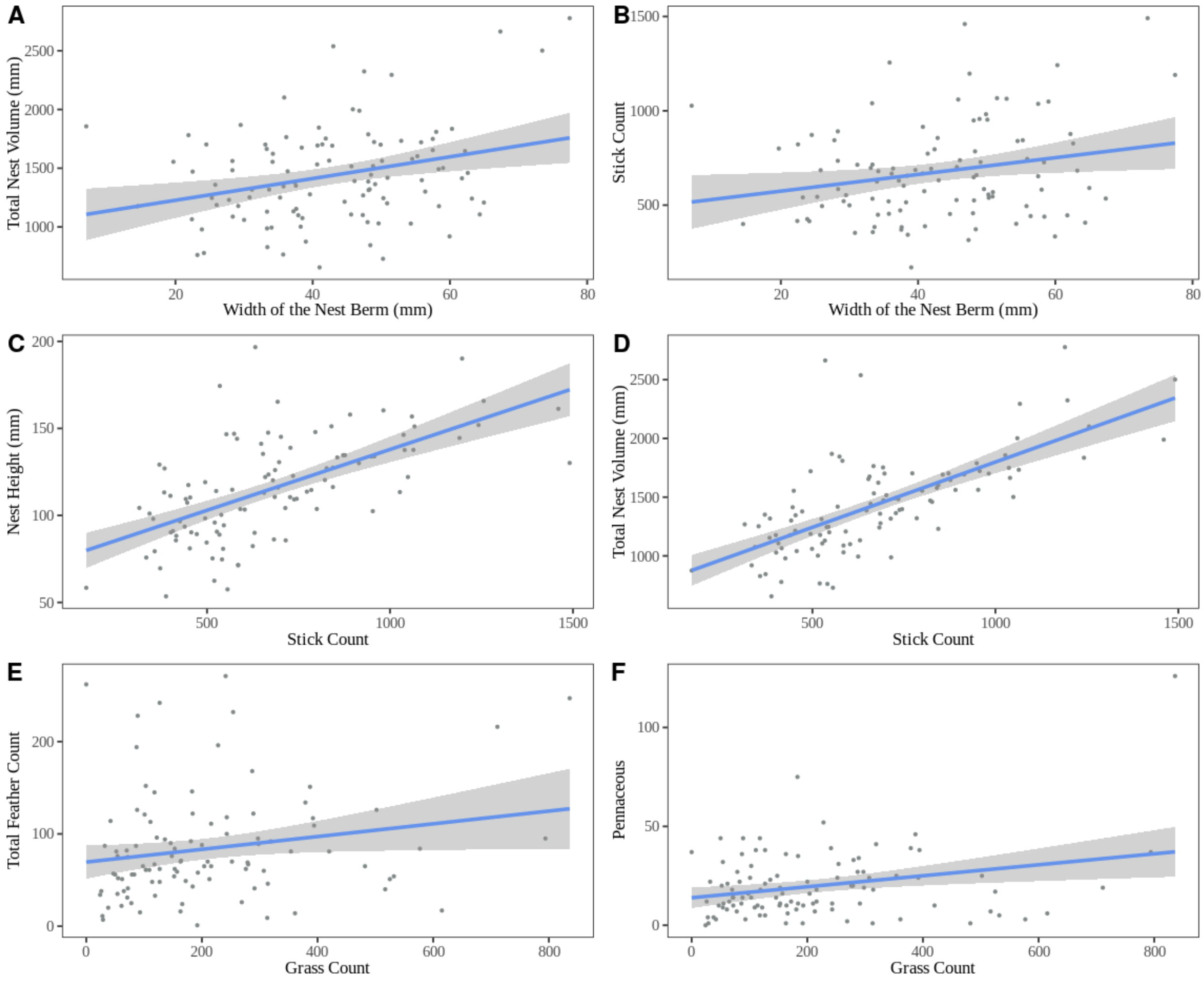


*Figure S2 - Relationships between different nest attributes and building materials. The width of the nest berm was positively correlated with the total volume (A;* $r^2 = 0.09$, $P < 0.001$*) and stick count (B;* $r^2 = 0.04$, $P = 0.02$) *of the nest. The stick count further highly positively correlated with the height of the nest (C;* $r^2 = 0.37$, $P < 0.001$*) and the total volume of the nest (D;* $r^2 = 0.41$, $P < 0.001$*). The grass count positively correlated with the total count of feathers (E;* $r^2 =$

0.03, $P = 0.04$) *and pennaceous feather count (F;* $r^2 = 0.06$, $P = 0.005$)*. Gray shading encompasses the 95% confidence interval for the line of best fit.*

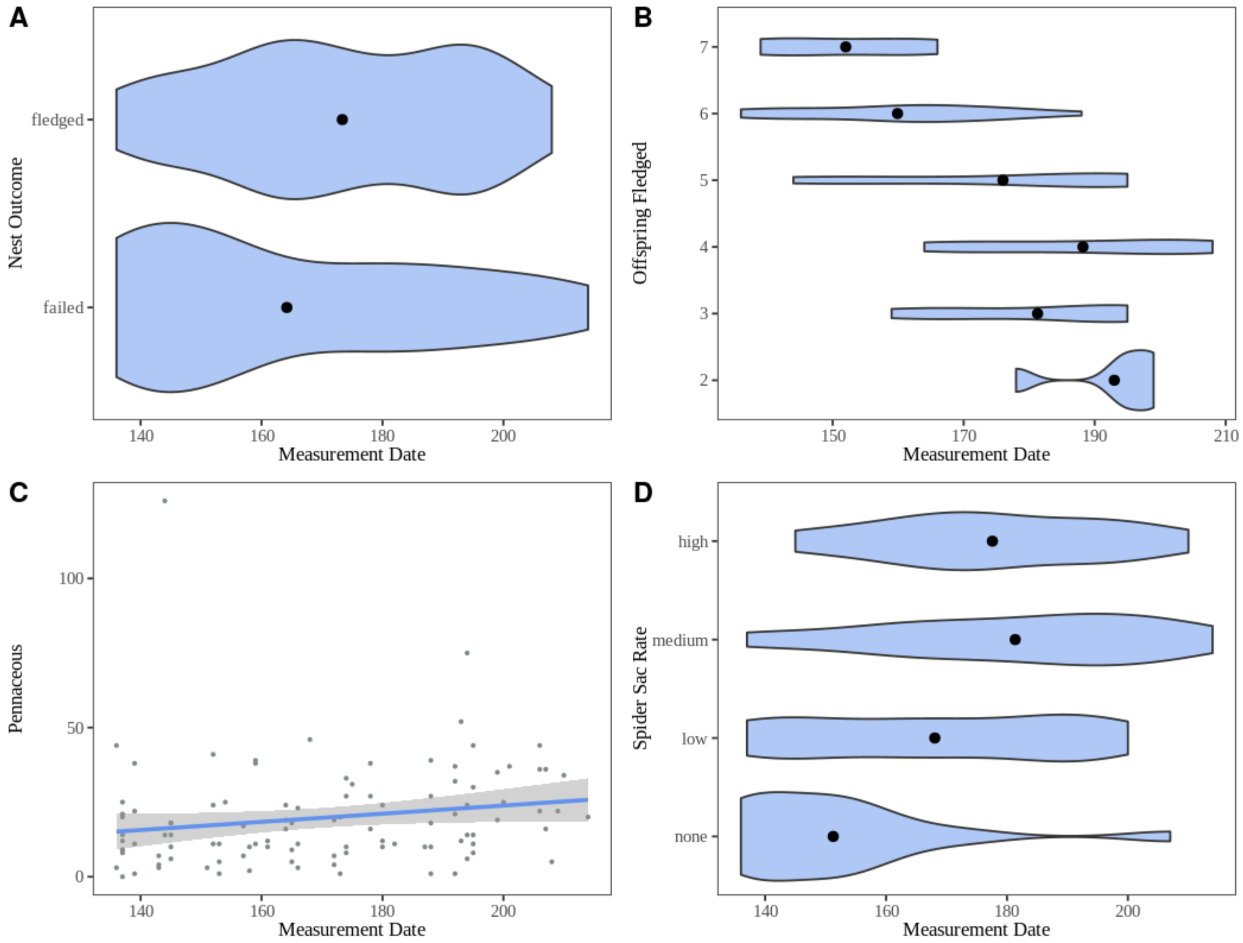


*Figure S3 - Significant relationships between nest characteristics and the date of nest measurement (i.e., completion of egg laying). Nests measured earlier in the season were more likely to fail than nests measured later in the season (A;* $t = 2.04$, *$P = 0.04$), but also fledged fewer offspring (B;* $t = -5.89$, *$P = 0.002$). The pennaceous feather count positively correlated with the measurement date after the removal of the outlier on DOY 144 (C;* $r^2 = 0.02$, *$P = 0.001$). Nests measured later in the season also contained more spider sac material than those*

*measured earlier in the season (D;* t = 4.49, $P < 0.001$*). Gray shading in C encompasses the 95% confidence interval for the line of best fit.*

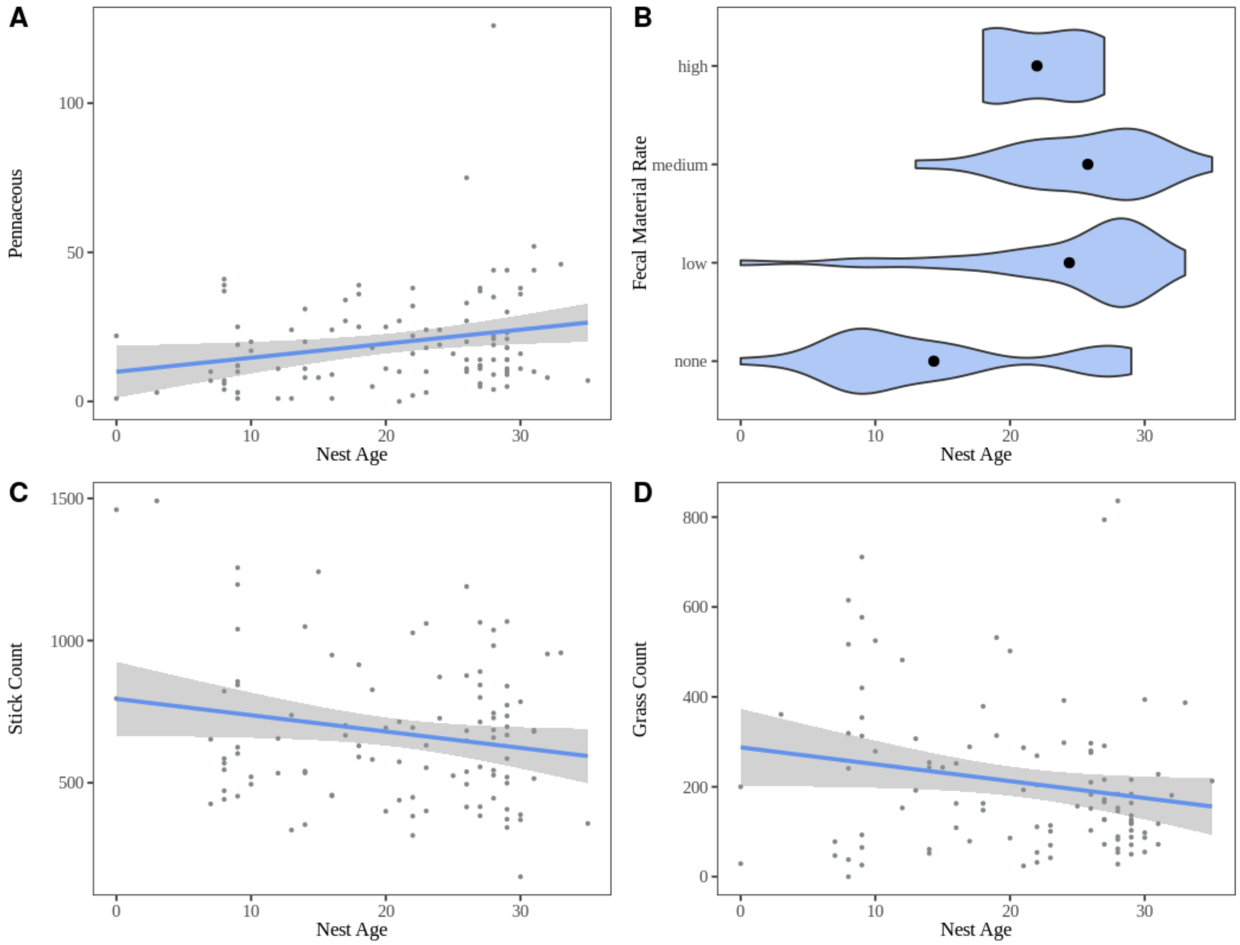


*Figure S4 - Relationship between three nest materials and fecal material with the age of the nest at collection. Pennaceous feather count positively correlated with the nest age (A;* $r^2 = 0.04$, $P = 0.02$*). Although not statistically significant, the stick count (C;* $r^2 = 0.03$, $P = 0.052$*) and the grass count (D;* $r^2 = 0.03$, $P = 0.054$*) negatively correlated with the age of the nest. Older nests contained more fecal material than younger nests* (B; t = 5.24, $P < 0.001$). *Gray shading in A, C, and D encompasses the 95% confidence interval for the line of best fit.*
